# Maternal Piwi Regulates Primordial Germ Cell Development to Ensure the Fertility of Female Progeny in *Drosophila*

**DOI:** 10.1101/2021.04.30.442025

**Authors:** Lauren E Gonzalez, Xiongzhuo Tang, Haifan Lin

## Abstract

In many animals, germline development is initiated by proteins and RNAs that are expressed maternally. PIWI proteins and their associated small noncoding <u>P</u>IWI-<u>i</u>nteracting RNAs (piRNAs), which guide PIWI to target RNAs by base-pairing, are among the maternal components deposited into the germline of the early embryo in *Drosophila*. Piwi has been extensively studied in the adult ovary and testis, where it is required for transposon suppression, germline stem cell self-renewal, and fertility. Consequently, loss of Piwi in the adult ovary using *piwi*-null alleles or knockdown from early oogenesis results in complete sterility, limiting investigation into possible embryonic functions of maternal Piwi. In this study, we show that the maternal Piwi protein persists in the embryonic germline through gonad coalescence, suggesting that maternal Piwi can regulate germline development beyond early embryogenesis. Using a maternal knockdown strategy, we find that maternal Piwi is required for the fertility and normal gonad morphology of female, but not male, progeny. Following maternal *piwi* knockdown, transposons were mildly derepressed in the early embryo but were fully repressed in the ovaries of adult progeny. Furthermore, the maternal piRNA pool was diminished, reducing the capacity of the PIWI/piRNA complex to target zygotic genes during embryogenesis. Examination of embryonic germ cell proliferation and ovarian gene expression showed that the germline of female progeny was partially masculinized by maternal *piwi* knockdown. Our study reveals a novel role for maternal Piwi in the germline development of female progeny and suggests that the PIWI/piRNA pathway is involved in germline sex determination in *Drosophila*.

## INTRODUCTION

Maternally deposited proteins and mRNAs drive the earliest stages of embryonic development. This is achieved in a unique way during the germline development of many animals, including *Drosophila*. During *Drosophila* oogenesis, germline-determining maternal proteins and RNAs are enriched in the cytoplasm at the posterior of the developing oocyte, in a region called the germ plasm (Lehmann 2016). This concentration of germline determinants facilitates the rapid and local specification of posterior nuclei into primordial germ cells (PGCs) during the maternal phase of embryogenesis. PGCs remain largely transcriptionally quiescent for several hours after the zygotic transcriptome is activated in the soma (Siddiqui et al. 2012; Van Doren et al. 1998). Even after the activation of the zygotic genome in the germline, the migration of PGCs through the midgut and their coalescence into embryonic gonads is regulated by intrinsic maternal and zygotic factors, and by the surrounding somatic cells (Jemc 2011; Slaidina and Lehmann 2017; Wawersik et al. 2005).

*Drosophila* PIWI proteins – Piwi, Aubergine (Aub), and Argonaute3 (Ago3) – are among these germ plasm-enriched maternal factors (Brennecke et al. 2007; Harris and Macdonald 2001; Mani et al. 2014; Megosh et al. 2006). The vast majority of embryos laid by *piwi-*null, *aub*-null, and *ago3-*null females suffer chromosome segregation defects which result in embryonic arrest before gastrulation (Mani et al. 2014). Of the few embryos that progress beyond the first few cell cycles, those laid by *aub*-null and *piwi-*null females fail to specify PGCs (Harris and Macdonald 2001; Megosh et al. 2006). In the case of the embryos from *aub*-null females, this is likely because Aub is required for the enrichment of germ plasm mRNAs at the posterior of the embryo by stabilizing them in the posterior and destabilizing them in other regions (Barckmann et al. 2015; Rouget et al. 2010; Vourekas et al. 2016). Aub, like other PIWI proteins, can regulate such diverse RNAs because of their association with small noncoding <u>P</u>IWI-<u>i</u>nteracting RNAs (piRNAs) (Aravin et al. 2006; Brennecke et al. 2007; Girard et al. 2006; Grivna et al. 2006; Lau et al. 2006). *Drosophila* piRNAs are 23-30 nt long, extremely heterogeneous, and can bind target RNAs with imperfect complementarity (Gou et al. 2014; Halbach et al. 2020; Shen et al. 2018; Zhang et al. 2018). Thus, they can guide PIWI proteins to potentially regulate a large swath of the transcriptome (Barckmann et al. 2015; Lee et al. 2012; Vourekas et al. 2016). All three *Drosophila* PIWI proteins can target mRNAs in the cytoplasm and regulate them post-transcriptionally, but Piwi mainly targets RNAs in the nucleus and regulates them at the transcriptional level.

The PIWI/piRNA pathway has been mostly studied in adult gonads, where its best-known role is in preventing transposon activation and protecting germ cells from transposon-induced genome instability (Wang and Lin 2021). Investigation into the role of PIWI/piRNA in transposon silencing was initially motivated by the observation that the majority of *Drosophila* piRNAs are complementary to transposons (Saito et al. 2006; Vagin et al. 2006). Building on this knowledge, it has been proposed that one major function of maternal PIWI/piRNA deposition into the germ plasm is to maintain transposon suppression across generations. This is exemplified in hybrid dysgenesis, in which progeny that paternally inherit a transposon but do not maternally inherit the corresponding transposon-targeting piRNAs cannot suppress the expression of that transposon (Akkouche et al. 2013; Brennecke et al. 2008). Consequently, hybrid dysgenesis typically results in genome instability, gonadal atrophy, and infertility (Dorogova et al. 2017; Engels and Preston 1979; Kidwell et al. 1977). Many studies argue that developmental defects in PIWI/piRNA mutants are caused by transposon derepression, but some studies suggest that transposon derepression is separable from other defects (Durdevic et al. 2018; Klenov et al. 2011).

Piwi and other PIWI/piRNA pathway proteins have been observed in germ cells throughout embryogenesis (Marie et al. 2017), and the function of zygotic Piwi in embryogenesis has begun to be explored in the context of transposon suppression (Akkouche et al. 2017; Marie et al. 2017). Although most *Drosophila* piRNAs target transposons, there are also piRNAs that target and regulate non-transposon genes in the early embryo (Barckmann et al. 2018; Barckmann et al. 2015; Dufourt et al. 2017; Ramat et al. 2020; Rouget et al. 2010), adult ovaries (Klein et al. 2016; Lin et al. 2020; Ma et al. 2017; Rojas-Rios et al. 2017), and adult testes (Gonzalez et al. 2015; Kotov et al. 2019; Nishida et al. 2007). This raises the possibility that maternal Piwi/piRNA could regulate non-transposon genes beyond early embryogenesis. To date, studies into this possibility have largely been limited by the severe cell cycle defects during early embryogenesis (Durdevic et al. 2018; Mani et al. 2014) and failure of PGC specification (Megosh et al. 2006) in progeny of females with mutations in PIWI/piRNA pathway genes. This challenge is further complicated in the case of Piwi itself, because *piwi* mutations block germline stem cell self-renewal, so *piwi* mutant females do not complete oogenesis (Cox et al. 1998; Cox et al. 2000). Only genetically mosaic females with Piwi expressed in ovarian somatic but not germline cells can lay eggs that lack maternal Piwi (Cox et al. 1998; Cox et al. 2000; Mani et al. 2014; Megosh et al. 2006). However, most of these eggs arrest in early embryogenesis (Cox et al. 1998; Mani et al. 2014). Because of these technical limitations, it remains unexplored whether maternal Piwi regulates any developmental processes beyond early embryonic stages.

In this study, we investigated the function of maternal Piwi on the germline development of progeny. We first observed that maternal Piwi persists in the embryonic germline at least through gonad coalescence. We then used a UAS/Gal4 approach to reduce but not completely eliminate maternal Piwi levels, which circumvents the aforementioned challenges of studying the function of maternal Piwi. Strikingly, we found that knockdown of maternal *piwi* decreased the fertility of female, but not male, progeny, suggesting that maternal Piwi plays a critical role in the germline development of female progeny. In contrast to other PIWI/piRNA depletion approaches, transposon expression was only mildly affected by maternal *piwi* knockdown. Transcriptome analysis, piRNA analysis, and examination of embryonic germ cell proliferation suggest that the germline of female progeny is partially masculinized by maternal *piwi* knockdown. These observations add to the growing body of evidence that the PIWI/piRNA pathway can regulate developmental processes independently of its transposon suppression function.

## METHODS

### Fly strains and husbandry

All *Drosophila* stocks were raised on standard agar/molasses medium and raised at 25°C for experiments. For all crosses involving UAS constructs and GAL4 constructs, females carried the UAS constructs and males carried the GAL4 constructs. Two anti*-piwi* shRNA lines were used: “Piwi RNAi #1,” which is BDSC stock #37483 and targets *piwi* exon 3, and “Piwi RNAi #2,” which was generated in Dr. Ting Xie’s lab (called “THU”) and targets *piwi* exon 2. *Df(2L)ED761* and *zuc^HM27^* fly stocks were gifts from Dr. Trudi Schupbach, and the *Dfd-lacZ* fly stock was a gift from Dr. Mark van Doren. See Reagent Table for more details about these and other fly stocks.

### Fertility tests

For female fertility tests, virgin females were mated with two *w^1118^* males in individual vials for two days before beginning the fertility test. For each vial, the flies were transferred to a new vial every 24 hours, and the number of eggs in the old vial was counted within four hours of transferring the parents. If the female died during the fertility test, the day of death was recorded and fertility data were omitted. If a male died during the fertility test, it was replaced with another *w^1118^* male.

For male fertility tests, each vial contained one male with three virgin *w^1118^* females, vials were transferred every 24 hours, and the number of eggs in the old vial were counted within four hours of transferring the parents. On alternating days, the three females were replaced with three new virgin *w^1118^* females. If a female or male died during the fertility test, the day of death was recorded and fertility data were omitted.

#### For all fertility tests

Minimal yeast was added to vials to minimize nutrition-related effects on fertility and to aid in visualization of eggs. Each vial was allowed to age until pupae began eclosing. Once pupae began eclosing, the number of adults were counted every day until all were eclosed. Percentage of adults eclosed was calculated as (# total adults) / (# embryos). Males and females were counted separately, and sex ratios were calculated as (total # female adults) / (total # male adults). Percentage of adults eclosed and sex ratios were only calculated for parents who laid at least 10 eggs. 18-22 total individuals of each genotype were tested across two experiments performed at different times.

### Immunostaining

Ovaries from 2-3 day old females were dissected in 1× PBS, then fixed for 20 minutes in fixative (0.5% (v/v) NP-40, 2% formaldehyde in PBS) at room temperature. Ovaries were then washed in PBST (0.1% TritonX-100 in PBS) 3× for 15 minutes each, then blocked in 5% normal goat serum in PBST (NGS) for one hour at room temperature or overnight at 4°C. Blocking solution was replaced with primary antibody diluted in NGS, and samples incubated overnight at 4°C. Ovaries were again washed at room temperature in PBST 3× for 15 minutes each, and incubated in secondary antibody diluted in NGS for four hours at room temperature or overnight at 4°C. Ovaries were washed in PBST once, then incubated in DAPI diluted in PBST (1:5000) for 10 minutes and mounted in Vectashield.

Testes from 1-4 day old males were dissected in 1× PBS, then fixed in testis fixative (0.02% Triton-X-100, 2% formaldehyde in PBS) for 10 minutes, before being washed 3× for 15 minutes each in PBST (0.1% Triton-X-100). From this point onwards, testicular immunostaining proceeded as for ovaries above.

The third instar larval gonads were dissected in 1× PBS and fixed in fixative (100 mM glutamic acid, 25 mM KCI, 20 mM MgSO4, 4 mM 4 mM Sodium Phosphate, 1mM MgCI2 and 4% paraformaldehyde) at room temperature for 40 min, then briefly washed in PBST (PBS with 0.1% Triton X-100) and blocked with blocking buffer (PBST+0.5% normal goat serum) at room temperature for 30 min, followed by overnight incubation with primary antibody at 4°C. From this point onwards, immunostaining proceeded as described for ovaries above.

Embryos were collected on grape juice plates in embryo collection cages containing around 100 females and 20 males, washed in saline solution (0.12 M NaCl, 0.03% Triton-X-100), dechorionated in 50% bleach for two minutes, and washed thoroughly in deionized water. Embryos were then transferred to 50% heptane and 50% embryo fixation solution (1× PBS, 50 mM EGTA, 9.25% formaldehyde) and incubated for 10-20 minutes, rocking at room temperature. After removing fixative, embryos were washed in ice-cold 100% methanol (MeOH) and shaken for two minutes. Embryos were washed 3-4× in ice-cold 100% MeOH and either stored in MeOH at −20°C or immediately rehydrated in PBST by sequentially replacing MeOH with a MeOH:PBST series with increasing concentrations of PBST (5 minutes each in 70:30 MeOH:PBST, 50:50 MeOH:PBST, 30:70 MeOH:PBST, 100% PBST). Embryos were blocked in NGS for one hour at room temperature or overnight at 4°C, and immunostaining proceeded as described for ovaries above.

Samples were imaged using ZEISS Axio Imager2 or Leica TCS SP5 Spectral Confocal Microscope on sequential scanning mode, and image adjustments were made in ImageJ. See the Supplementary Reagent Table for details about antibodies and other commercial reagents.

### Western blotting

0-1.5 h embryos were collected on grape juice plates in embryo collection cages containing about 100 females and 20 males. Embryos were washed with deionized water, dechorionated in 50% bleach for 1 minute, washed again thoroughly with water, and transferred to MCB buffer (50 mM Hepes/NaOH (pH 7.5), 150 mM potassium acetate, 2 mM magnesium acetate, 10% (v/v) glycerol, 1 mM DTT, 0.1% (v/v) Triton X-100, 0.1% (v/v) NP-40, 1× protease inhibitor and 0.5% (v/v) beta-mercaptoethanol). Sample was homogenized with a pestle, then centrifuged at 14000 g for 20 minutes (4°C), and supernatant was transferred to a new tube. Approximately 50 μg of protein sample, heat denatured with 6× SDS loading dye, was run on a 10% SDS-PAGE gel. Proteins were transferred to a PVDF membrane, incubated with the appropriate primary antibody at 4°C overnight, then detected with the appropriate HRP-labeled secondary antibody. See the Supplementary Reagent Table for details about antibodies and other commercial reagents.

### RNA extraction and RT-PCR

Ovaries were extracted from 1-2 day-old females in 1× PBS, then transferred to TRIzol; RNA was extracted following manufacturer’s instructions. 6-8 pairs of ovaries were grouped together for each sample. 0-1.5 h embryos were collected as described for Western Blotting above and transferred to 1× PBST. PBST was replaced with TRIzol; RNA was extracted following manufacturer’s instructions.

Two μg of total RNA was used for reverse transcription using MultiScribe (ThermoFischer) following manufacturer’s instructions, and diluted 5× before quantitative PCR. For quantitative PCR, transcripts of interest were amplified using iTaq Universal SYBR Green (BioRad) using manufacturer’s recommended conditions, on a BioRad CFX96 Real-Time machine. All primer sets used for qPCR were confirmed to have amplification efficiency 80-110%. Gene expression was calculated using the ΔΔC_T_ method, using *act5C* as the housekeeping gene and using *GFP-MatKD* as the control, unless otherwise specified. For non-quantitative PCR, genes of interest were amplified using GoTaq Green Master Mix (Promega) with the following reaction conditions: 95°C 120s, 25× (95°C 30s, 55°C 30s, 72°C 60s), 72°C 300s. See the Supplementary Reagent Table for details about primers and other commercial reagents.

### Small RNA extraction, library preparation, and sequencing

For Small RNA-Seq, 20 μg of total RNA from 0-1.5 h embryos was run for 3.5 hours on a 15% polyacrylamide gel that had been pre-running in 1× TBE for at least one hour. RNA was visualized by incubating the gel in Diamond Dye (Promega) and the 20-29 nt area of the gel was cut out for purification. 20 nt / 30 nt / 70 nt RNA ladder was custom made. RNA was purified by fragmenting the gel piece and incubating fragments in AES buffer (300 mM NaAc, 2 mM EDTA 0.1% (w/v) SDS) overnight at room temperature. Supernatant was transferred to precipitation buffer (0.3 M NaAc, 90% ethanol (EtOH), 1 μl glycoblue (ThermoFischer) and incubated on ice for one hour. RNA was then centrifuged at 13000 g for 30 minutes, washed in 70% EtOH, resuspended in 10 μL nuclease-free H_2_O, and stored at −80°C. 5 μL (approximately 75 ng RNA) was used for library preparation.

Total RNA purified by TRIzol was used for library preparation for RNA-Seq. For both total RNA-Seq and Small RNA-Seq, three replicates of each sample were collected and analyzed. Sequencing libraries were prepared according to the Illumina TruSeq Small RNA Library Prep Kit or Illumina TruSeq Stranded mRNA Library prep protocols. All sequencing data were generated on a HiSeq2000 at the Yale Stem Cell Center Genomics Core and sequencing quality was assessed using FastQC (Andrews 2010).

### Bioinformatics analysis

For Total RNA-Seq: reads were mapped to *D. melanogaster* Berkeley Drosophila Genome Project (BDGP) release 6 using STAR (Dobin et al. 2013), and counted at transposable element and gene features, defined by gene annotation in BDGP6.28, using TETools (Lerat et al. 2017). Expression level changes between experimental and control samples were calculated using DESeq2 (Anders and Huber 2010) with standard parameters. For each RNAi line, genes which were differentially expressed >1.5-fold (compared to *GFP-MatKD*) with a significance p-adjusted <0.05 were considered to be differentially expressed. Gene Ontology analysis was performed as a PANTHER Overrepresentation Test using GO database released 2020-03-23 (doi: 10.5281/zenodo.3727280).

For Small RNA-Seq: Adapter sequence (TGGAATTCTCGGGTGCCAAGG) was trimmed from reads using CutAdapt. rRNA, miRNA, tRNA, and siRNA (according to FlyBase annotation, and including exogenous siRNAs being expressed by our RNAi lines: TTGTAGTTGTACTCCAGCTTG for GFP RNAi, TTGATTTCGGAGTTTGTCCAA for Piwi RNAi #1, and TTCGTCTGCAGCATGACCGGGG for Piwi RNAi #2) were filtered out, and remaining sequences were then mapped to the *Drosophila* genome using Bowtie1.2.2 (Langmead et al. 2009) to create the list of putative piRNAs. Nucleotide composition was determined using FastQC (Andrews 2010). To identify putative piRNA targets on non-transposon mRNAs, these putative piRNAs were mapped using Bowtie1.2.2 (Langmead et al. 2009) to the transcribed sequences (BDGP sequence release 6, annotation release 25; dmel-all-gene-r6.26.fasta retrieved from FlyBase) of all *D. melanogastar* genes, excluding transposons. Mapping required the piRNA read to be antisense to the mRNA and allowed up to 2 mismatches in the first 24 nucleotides. Featurecounts (Liao et al. 2014) was used to count the number of piRNA reads that aligned to each mRNA, weighted for the number of mapping locations for each read. This count is referred to as “piRNA targetability.” Targetability level changes between experimental and control samples was calculated using DESeq2 (Anders and Huber 2010) with standard parameters. For each RNAi line, genes which were differentially targetable >1.5-fold (compared to *GFP-MatKD*) with a significance p-adjusted <0.05 were considered to be differentially targetable. Full scripts for piRNA target analysis available in the Supplemental Materials.

### Data availability

Fly lines and antibodies are available from their original commercial or lab source. Anti-Piwi 4K5 antibody is available from the Haifan Lin lab upon request. RNA-seq and Small RNA-seq sequence data are available at the National Center for Biotechnology Information GEO database with the accession number GSE171951.

## RESULTS

### Maternal Piwi persists in germ cells throughout embryogenesis

To begin investigating the function of maternal Piwi, we first characterized its expression pattern in embryos using the cross depicted in Fig 1A. Transheterozygous *myc-piwi/UASp-GFP* (3^rd^ chromosome) females were crossed with homozygous *zfh2-gal4* (3^rd^ chromosome) males that express GAL4 throughout the nervous system (Jenett et al. 2012; Lai et al. 1991). The *UASp-GFP/zfh2-gal4* progeny inherit maternal Myc-Piwi protein and express GFP but do not express Myc-Piwi from the zygotic genome. In contrast, *myc-piwi/zfh2-gal4* progeny inherit maternal Myc-Piwi and express zygotic Myc-Piwi, but not GFP. With this approach, we were able to definitively identify *UASp-GFP/zfh2-gal4* embryos and use anti-Myc immunostaining to visualize maternal Piwi persistence during embryogenesis.

**Figure 1.**
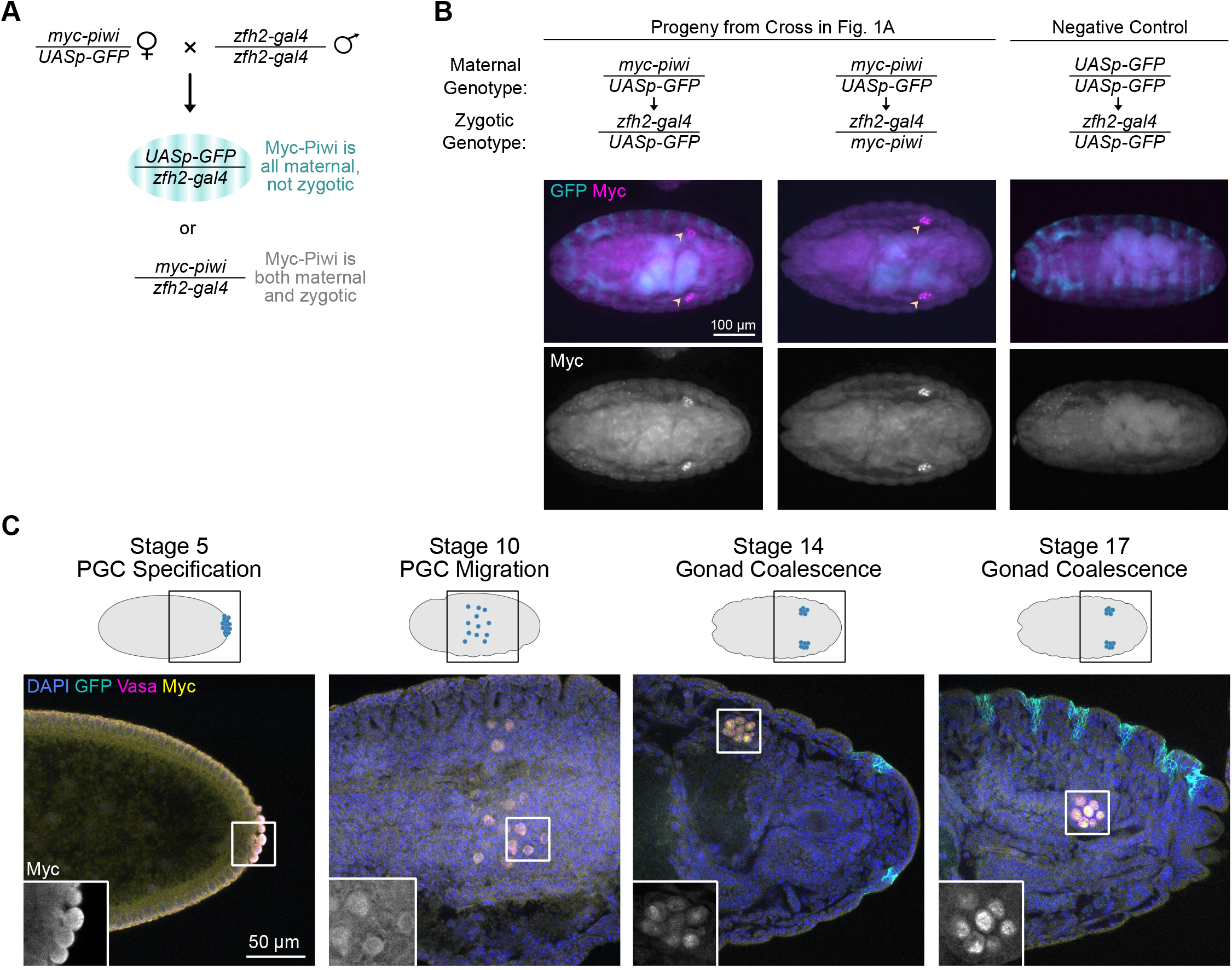
Maternal Piwi persists in germ cells through gonad coalescence. (A) Schematic of crossing strategy to visualize maternal Piwi. (B) Representative images from visualization of Myc-Piwi in late-stage embryos (approximately 16 hours after egg laying) inheriting only maternal Myc-Piwi protein (left panel), inheriting both maternal Myc-Piwi protein and the *myc-piwi* transgene (middle panel), and inheriting neither the Myc-Piwi protein nor the *myc-piwi* transgene (right panel; negative control). Coalesced gonads indicated by arrowheads. (C) Representative images of maternal Myc-Piwi protein in *UASp-GFP/zfh2-gal4* embryos from the cross depicted in Fig 1A, and schematics of germ cell localization at each developmental timepoint. Inset is magnified 2X. PGC: primordial germ cells.

We observed strong maternal Myc-Piwi expression in *UASp-GFP/zfh2-gal4* embryos up to at least 17 hours after egg laying (Fig 1B, left panels). At these relatively late stages of embryogenesis, maternal Myc-Piwi was detectable in a number of cell types, but most strongly in primordial germ cells (PGCs) (Fig 1B, left panels). As expected, Myc-Piwi was also strongly detectable in the PGCs of *myc-piwi/zfh2-gal4* embryos, in which Myc-Piwi represents a mixture of maternal and zygotic Piwi (Fig 1B, middle panels). From PGC specification through migration, maternal Myc-Piwi localized within both the cytoplasm and nucleus, as previously described for maternal Piwi in early PGCs (Mani et al. 2014), but become predominantly nuclear in coalesced gonads (Fig 1C). These observations suggest that maternal Piwi could have a direct role in germline development throughout embryogenesis, including germline processes that occur during or after gonad coalescence.

To visualize maternal Piwi in the larval gonad, we used a similar strategy, this time crossing transheterozygous *myc-piwi/UASp-GFP* females to *Act-gal4* males (Fig S1). In *UASp-GFP/Act-gal4* third instar larvae in which Myc-Piwi represents only maternal Piwi, all testes and most ovaries lacked detectable maternal Myc-Piwi, but about 20% of ovaries did contain detectable maternal Myc-Piwi (Fig S1). This suggests that maternal Piwi is present during some stages of female larval gonadogenesis, up to the third instar stage.

### Knockdown of maternal *piwi* resulted in female-specific infertility in the F1 generation

To assess the role of maternal Piwi in the germline development of progeny, we used *Maternal Alpha Tubulin (MAT)-gal4* and two independent anti-*piwi* shRNA lines to reduce the levels of maternal Piwi (herein referred to as *piwi-MatKD* #1 and #2; Fig 2A). In these females (F0 females in Fig 2A), Piwi is knocked down specifically in germ cells starting in mid-oogenesis (Fig 2B). In contrast to strong *piwi-*null mutations where *piwi* expression in all ovarian cells is depleted, knocking down *piwi* expression in the germline did not affect germline stem cell self-renewal and resulted in F0 ovaries with grossly normal morphology (Fig 2B). When *piwi-MatKD* F0 females are crossed with *w^1118^* males, half of the F1 progeny carries a copy of *UASp-piwi-shRNA* and the other half carries a copy of *MAT-gal4* (Fig 2A). *MatKD* F1 flies used throughout this study were a mixture of these two genotypes. Although the maternal deposition of GAL4 protein could in theory activate the shRNA in the half of F1 progeny that carry *UASp-shRNA* (Staller et al. 2013), we saw barely detectable levels of zygotic Piwi protein and RNA during early- and mid-embryogenesis (Fig S2), so even if *piwi-*shRNA could reduce zygotically expressed *piwi*, the overall impact of this reduction as compared to the knockdown of maternal Piwi would be negligible.

**Figure 2.**
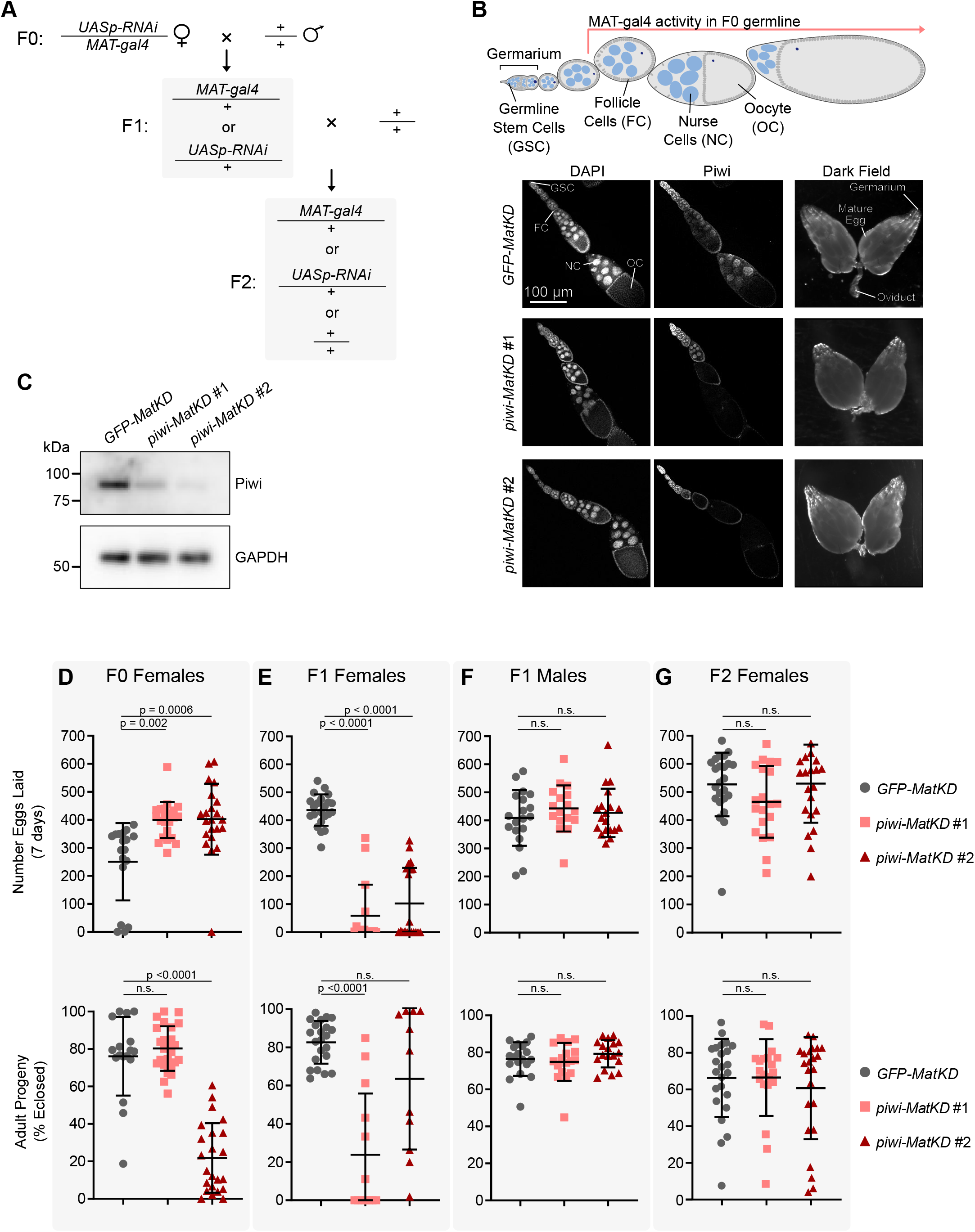
Maternal Piwi is required for fertility of female, but not male, progeny. (A) Schematic of crossing strategy to deplete maternal Piwi. F0 individuals carry both *UASp-shRNA* and *Maternal Alpha Tubulin (MAT)-GAL4*, so shRNA is expressed from mid-oogenesis. F1 individuals carry either *UASp-shRNA* or *MAT-GAL4*, but not both. (B) Schematic of MAT-GAL4 expression in the *MatKD* F0 ovary, and anti-Piwi immunofluorescence in *MatKD* F0 ovaries. Whole ovary images are at the same scale. (C) Western blot for total Piwi and total GAPDH in 0-2 h F1 embryos. (D-G) Seven-day fertility tests of individual females mated to two *w^1118^* males or individual males mated to three *w^1118^* females (see cross in Fig 2A). (D) *MatKD* F0 females (n=17-25). (E) *MatKD* F1 females (n=16-22). (F) *MatKD* F1 males (n=17-20). (G) *MatKD* F2 females (n=20-24). Upper panels indicate number of eggs laid per day and lower panels indicate the percentage of those eggs that developed to adulthood (only calculated for crosses which produced ≥ 10 eggs). Mean + SD. One-way ANOVA and Dunnet’s multiple comparisons test. n.s. = “not significant.”

Both anti-*piwi* shRNA lines efficiently reduced Piwi protein levels, but to different degrees: *piwi-MatKD* #1 was relatively weaker than *piwi-MatKD* #2 (Fig 2C). Accordingly, embryos laid by *piwi-MatKD* #1 (“*piwi MatKD* #1 F1” embryos) developed largely normally, whereas *piwi-MatKD* #2 F1 embryos frequently arrested at pre-cellularization stages (Fig S3). This *piwi-MatKD* #2 F1 phenotype is reminiscent of the F1 embryonic arrest that results from maternal Piwi depletion using strong *piwi*-null ovarian clones (Mani et al. 2014). Despite the high rate of embryonic arrest, some *piwi-MatKD* #2 F1 embryos did progress to adulthood (see below).

We performed fertility tests of *piwi*-*MatKD* F0, F1, and F2 flies by crossing them individually with 2-3 *w^1118^* flies of the reciprocal sex, counting the number of eggs laid over seven days and the total number of adults that emerged from these vials after 12-15 days (Fig 2D-G). In contrast to *piwi-*null females, *piwi-MatKD* F0 females had normal ovary morphology (Fig 2B) and normal egg-laying capacity (Fig 2D, upper panel). In fact, they laid more eggs than control *GFP-MatKD* F0 females, possibly because the *GFP-MatKD* F0 females were somewhat compromised in egg-laying (Fig 2D, upper panel). Up to 100% of *piwi-MatKD* #1 F1 embryos and up to 60% of *piwi-MatKD* #2 F1 progressed to adulthood (Fig 2D, lower panel), consistent with the incomplete penetrance of embryonic arrest in *piwi-MatKD* F1 #2 embryos (Fig S3). These F1 adults provided us with an opportunity to observe additional developmental defects in *piwi-MatKD* F1 flies.

Because we had observed that maternal Piwi was specifically maintained in the F1 embryonic germline (Fig 1), our measurement of the fertility of *piwi-MatKD* F1 females allowed us to assess whether maternal Piwi had any effect on progeny germline development. Indeed, *piwi-MatKD* F1 females from both anti-*piwi* shRNA lines suffered dramatic fertility defects (Fig 2E, Fig S4). This was reflected in both decreased rates of egg-laying (Fig 2E, upper panel) and decreased percentages of those eggs developing to adulthood (Fig 2E, lower panel), as compared to *GFP-MatKD* F1 controls. Notably, although approximately 60% of *piwi-MatKD* #2 F1 embryos could not progress beyond the first few cell cycles of embryogenesis (Fig S3), *piwi-MatKD* #2 F1 adult females were equally subfertile as *piwi-MatKD* #1 F1 females (Fig 2E).

If the *piwi-MatKD* F1 subfertility was caused by some general germline development defect, such as failure to specify PGCs during embryogenesis (Megosh et al. 2006), the fertility of both females and males would be affected. To our surprise, *piwi-MatKD* F1 males, when crossed with virgin *w^1118^* females, were able to stimulate egg laying (Fig 2F, upper panel) and fertilize those eggs (Fig 2F, lower panel) at rates indistinguishable from those of *GFP-MatKD* F1 males. The normal fertility of *piwi-MatKD* F1 males, together with the subfertility of *piwi-MatKD* F1 females, suggests that maternal Piwi plays some role in female-specific, rather than general, aspects of germline development in progeny.

We did not observe any subfertility in females in the *piwi-MatKD* F2 generation (Fig 2G, Fig S4, Fig S5), suggesting that zygotic *piwi* expression in the *piwi-MatKD* F1 ovary provides sufficient maternal Piwi for the normal germline development of *piwi-MatKD* F2 females. However, these F2 females were the progeny of the more mildly affected (not fully sterile) F1 females, and it is impossible to examine the phenotype of the progeny of severely affected (fully sterile) F1 females. Because of this, our measurement of F2 fertility is biased towards mildly affected females, and it remains unknown whether the defects of *piwi-MatKD* F1 females would be transmitted to their progeny.

To investigate what ovarian defects led to the subfertility of *piwi-MatKD* F1 females, we examined *MatKD* F1 gonads. Approximately 40% of *piwi-MatKD* F1 ovaries lacked late-stage egg chambers (“Arrested”) (Fig 3A, 3B). In these ovaries, oogenesis arrested at the onset of vitellogenesis (Fig 3B), a checkpoint where oogenesis can be stalled by a variety of stimuli ranging from environmental stress to imbalances in ecdysone or Juvenile Hormone levels (McCall 2004). Furthermore, 2-7% of *piwi-MatKD* F1 ovaries completely lacked germline cells (“Agametic”) (Fig 3A, 3B), a phenotype that evokes *piwi*-null mutant ovaries (Cox et al. 1998). The remaining 51-58% of *piwi-MatKD* F1 ovaries were morphologically wildtype-like (Fig 3A, 3B). These females with wildtype-like ovaries were nevertheless subfertile in comparison to *GFP-MatKD* F1 females (Fig S6), so the ovarian arrest was not the sole cause of *piwi-MatKD* F1 subfertility. All *piwi-MatKD* F1 ovaries had normal Piwi expression in both somatic and germline cells in non-degenerating egg chambers (Fig 3B), indicating that zygotic Piwi expression was not affected in *piwi-MatKD* F1 females.

**Figure 3.**
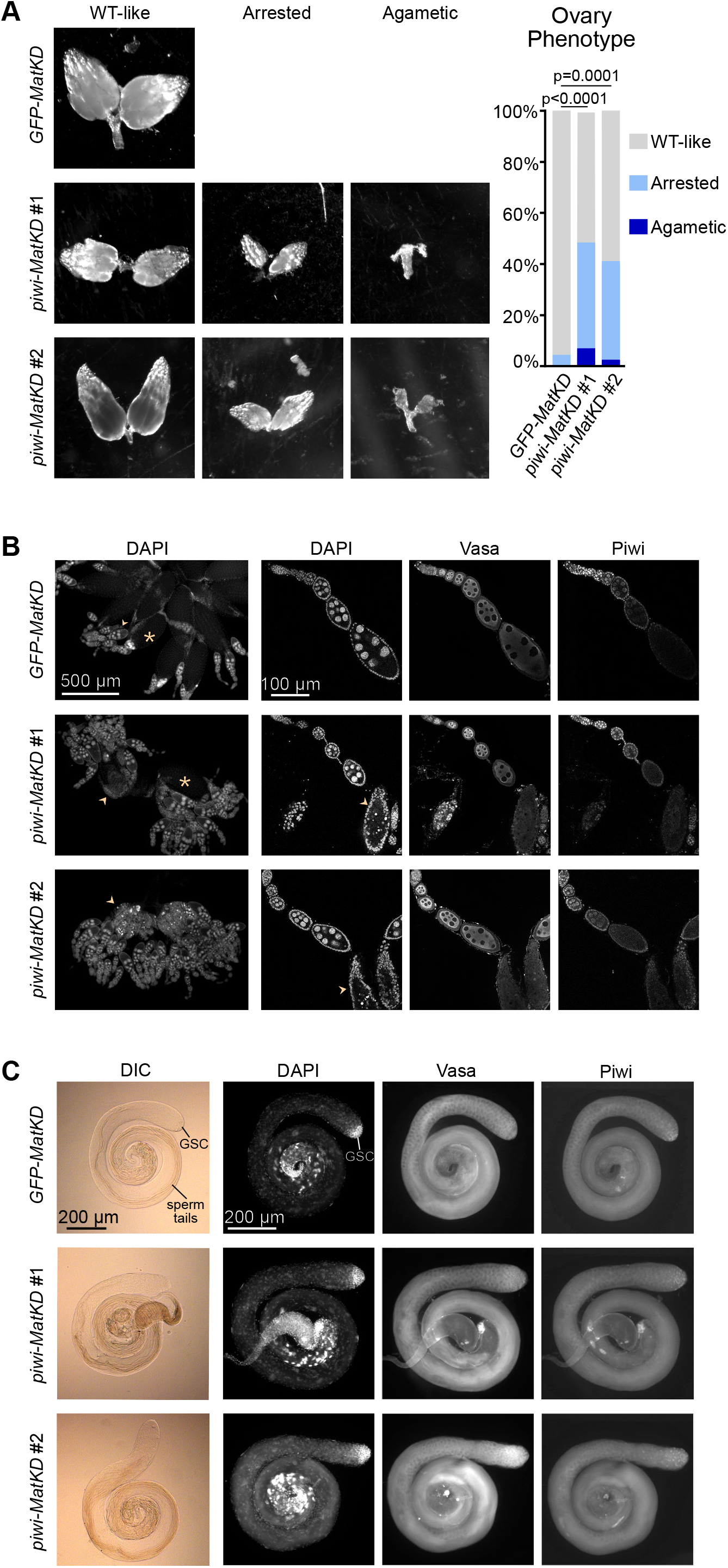
Adult females depleted of maternal Piwi had arrested ovaries. (A) Representative images of *MatKD* F1 ovaries (2-3 days old), and relative frequency of each category of ovary phenotype in each genotype (n=45-75 per genotype). Chi-square test. Whole ovary images are at the same scale. (B) Anti-Vasa and anti-Piwi immunofluorescence of *MatKD* arrested ovaries. *Piwi-MatKD* arrested ovaries arrest around stage 8. An asterisk indicates one example mature egg, and an arrowhead indicates one example arrested egg chamber. (C) Representative images of *MatKD* F1 testes, with anti-Vasa and anti-Piwi immunofluorescence, and Differentially Interference Contrast (DIC) to visualize sperm tails.

The testes of *piwi-MatKD* F1 males had no observable defects (Fig 3C), which further reinforces the female-specific nature of the F1 fertility defect.

### Transposons are mildly derepressed in *piwi-MatKD* F1 embryos and fully repressed in *piwi-MatKD* F1 adult ovaries

Given that the best-studied role of Piwi is in transposon repression, we next examined whether the *piwi-MatKD* F1 subfertility was a result of transposon derepression in the F1 embryo or the F1 adult ovary. To do so, we began by performing total RNA-seq on *piwi-MatKD* F1 0-1.5 h embryos (Fig S7, Table S1) and quantifying the expression of annotated transposons (Lerat et al. 2017). Only 11 transposons were derepressed under both *piwi-MatKD* conditions, with 16 derepressed in *piwi-MatKD* #1 F1 and 34 derepressed in *piwi-MatKD* #2 F1 embryos (Fig 4A, 4B). Among the derepressed transposons, retrotransposons in the LTR and LINE families were over-represented (Fig 4C). The expression of most of these derepressed transposons remained at relatively low levels, with the exception of *Het-A* (Fig S8), as compared to the typical derepression by 1-2 orders of magnitude for dozens of transposons when the Piwi/piRNA function is strongly disrupted (Rozhkov et al. 2013; Senti et al. 2015; Sytnikova et al. 2014).

**Figure 4.**
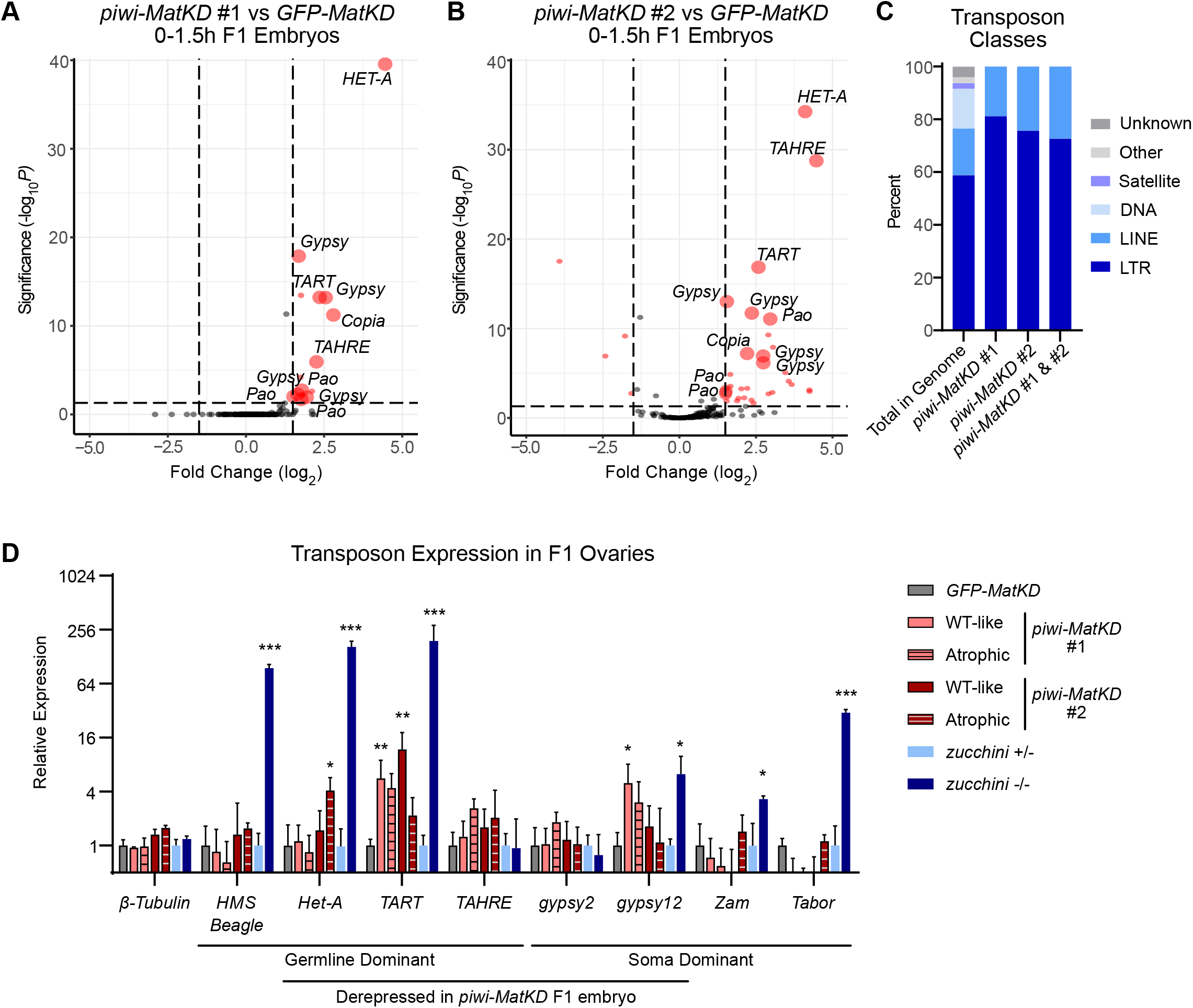
Transposons were marginally derepressed in the *piwi-MatKD* F1 generation. (A-B) Differential expression of transposons from RNA-seq on 0-1.5 h *MatKD* F1 embryos. Red datapoints are significantly changed (p-adjusted < 0.05, Fold Change > 1.5) in that genotype compared to *GFP-MatKD*, and large red datapoints are significantly changed (p-adjusted < 0.05, Fold Change > 1.5) in both *piwi-MatKD* #1 and #2 compared to *GFP-MatKD*. (C) Relative proportion of transposon classes in each group. (D) RT-qPCR for transposons in *MatKD* F1 adult ovaries, *zuc*^+/−^ (*zucchini^+/HM27^),* and *zuc*^−/−^ *(zucchini^HM27/Df(2L)716^*) ovaries. All *piwi-MatKD* groups were compared to *GFP-MatKD*, and *zuc*^−/−^ was compared to *zuc*^+/−^. The latter comparison serves as a positive control for transposon derepression in the context of piRNA pathway disruption. Two-way ANOVA and Dunnett’s Multiple Comparisons Test, * p<0.05, ** p<0.01, *** p<0.0001.

To investigate whether this moderate transposon derepression in the early embryo could seed further transposon derepression later in development, we used RT-qPCR to assess transposon expression in *MatKD* F1 ovaries. We separated the wildtype-like and arrested ovaries so we could confidently compare gene expression in ovaries of similar cellular composition (*GFP-MatKD* F1 and *piwi-MatKD* F1 with wildtype-like ovaries) while also identifying potential expression differences between wildtype-like and arrested ovaries of the same maternal genotype. We observed no consistent derepression of the transposons that had been moderately derepressed in the early *piwi-MatKD* F1 embryo (*Het-A*, *gypsy2, gypsy12, TART,* and *TAHRE*) nor of those that were derepressed upon zygotic Piwi knockdown during embryogenesis (*Het-A* and *HMS Beagle*) as previously reported (Akkouche et al. 2017) (Fig 4D). In a few cases (*Het-A, TART, gypsy12*), there was mild derepression in some but not all the *piwi-MatKD* groups, but their expression was still much lower than in *zuc*-null ovaries, where the piRNA pathway is truly disrupted. Thus, despite the mild transposon derepression in the early *piwi-MatKD* F1 embryo, their repression was restored by adulthood. These analyses of transposon RNA levels in embryo and adult stages suggest that the *piwi-MatKD* F1 subfertility was not mainly caused by transposon derepression and led us to explore whether it could instead have been caused by the mis-regulation of non-transposon genes.

### The maternally deposited mRNA and piRNA pools are shifted by maternal *piwi* knockdown

There is growing evidence that the PIWI/piRNA pathway also targets and regulates the expression of non-transposon genes (Wang and Lin 2021). As such, we used our RNA-seq of early F1 embryos to investigate whether *piwi-MatKD* had affected the broader maternally deposited transcriptome. Many more genes were differentially expressed in *piwi-MatKD* #2 than in *piwi-MatKD* #1 (Fig 5A). Many of the *piwi-MatKD* #2 differentially expressed genes function in global proteolysis and apoptosis (Table S2), which likely reflects the increased propensity of *piwi-MatKD* #2 F1 embryos to experience cell cycle defects and embryonic arrest (Fig S3) (Utz and Anderson 2000). Among the genes that were differentially expressed in both *piwi-MatKD* #1 and #2, 10 (including *piwi*) were down-regulated and 11 were up-regulated (Fig 5A, Table S3). As with all our analyses, we focused our attention on these gene expression changes in common between *piwi-MatKD* #1 and *piwi-MatKD* #2, because these are the most likely to explain the F1 subfertility that is in common between these two knockdown lines. Of the 21 genes commonly differentially expressed in the *piwi-MatKD* early embryo (Table S3), none was clearly related to the female-specific F1 subfertility, with the possible exception of *Jheh3* (see below).

**Figure 5.**
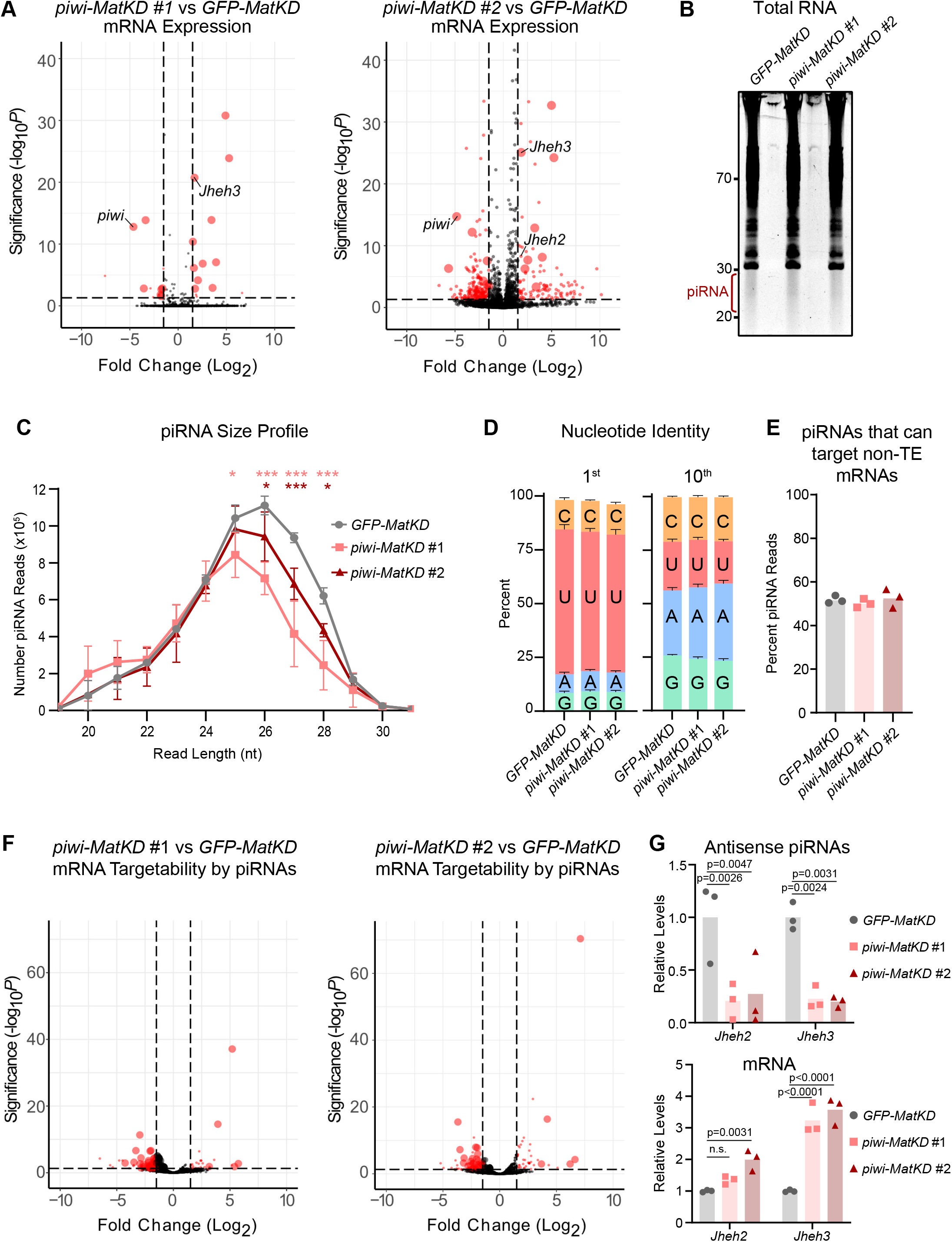
Knockdown of maternal *piwi* shifts the maternally deposited transcriptome and piRNA pool in the early F1 embryo. (A) Differential expression of non-transposon genes from RNA-seq on 0-1.5 h *MatKD* F1 embryos. Red datapoints are significantly changed (p-adjusted < 0.05, Fold Change > 1.5) in that genotype compared to *GFP-MatKD*, and large red datapoints are significantly changed (p-adjusted < 0.05, Fold Change > 1.5) in both *piwi-MatKD* #1 and #2 compared to *GFP-MatKD*. (B) Small RNAs (20 - 29 nt) were isolated from total 0-1.5 h *MatKD* F1 embryo RNA for Small RNA-seq. (C) Size distribution of piRNAs from total Small RNA-seq, after filtering out rRNA, miRNA, and siRNA, normalized to total library size. Two-Way ANOVA, Dunnett’s Multiple Comparisons Test. * p<0.01, ***p<0.0001. (D) Sequence distribution from each piRNA library at the first and tenth nucleotide position. (E) Percentage of reads in each piRNA library which have the capacity to target mRNAs, defined as being antisense to a transcribed gene region, allowing up to two mismatches. (F) Differential Targetability of *Drosophila* non-transposon mRNAs in *piwi-MatKD* vs *GFP-MatKD* F1 embryos. Red datapoints are significantly changed (p-adjusted < 0.05, Fold Change > 1.5) in that genotype compared to *GFP-MatKD*, and large red datapoints are significantly changed (p-adjusted < 0.05, Fold Change > 1.5) in both *piwi-MatKD* #1 and #2 compared to *GFP-MatKD*. (G) Putative antisense piRNA levels (upper panel, from Small RNA-seq) and mRNA levels (lower panel, from RNA-seq) for *Jheh2* and *Jheh3*. Relative levels determined from DESeq2. Significance tested with Two-Way ANOVA and Dunnett’s Multiple Comparisons Test.

Because we had observed maternal Piwi expression in the germline throughout embryogenesis (Fig 1), we next asked whether *piwi-MatKD* might cause the F1 infertility by dysregulating PIWI/piRNA-targeted genes in later developmental stages. To begin probing this possibility, we assessed the maternally deposited piRNA pool in *MatKD* F1 early embryos, reasoning that loss or accumulation of particular piRNAs in the early embryo could shape subsequent Piwi-mediated gene regulation throughout embryogenesis. We sequenced small RNAs isolated from total RNA of 0-1.5 h embryos (Fig 5B, Table S4) and found that the piRNA pool was reduced, though not abolished, by *piwi-MatKD* (Fig 5C). There was a particular reduction of 26-29 nucleotide piRNAs (Fig 5C), which are preferentially bound by Piwi rather than Aub or Ago3 (Brennecke et al. 2007). The remaining piRNAs in *piwi-MatKD* F1 embryos still largely retained the typical 1U and 10A biases, though the former was slightly diminished while the latter was slightly enhanced compared to piRNAs from *GFP-MatKD* F1 embryos (Fig 5D). These changes in the size distribution, 1U bias, and 10A bias suggest a shift from primary piRNAs to secondary piRNAs, which is consistent with a selective decrease of Piwi-bound piRNAs in the *piwi-MatKD* F1 early embryo.

To explore the possibility that these changes in the maternally deposited piRNA pool could impact PIWI/piRNA target genes later in development, we predicted target genes of the maternally deposited piRNAs. To guide PIWI proteins to a particular target, a piRNA must be able to base-pair with that RNA, though it does not need to base-pair perfectly (Barckmann et al. 2015; Gou et al. 2014; Halbach et al. 2020; Shen et al. 2018; Zhang et al. 2018). To identify target mRNAs, piRNA reads were aligned antisense to transcribed regions of non-transposon genes with up to two mismatches allowed. Each alignment was then weighted by the number of putative target sites of that piRNA read within the transcriptome. We refer to this calculated number of piRNA target sites within a gene as the “piRNA targetability” of that gene (see Methods and Supplemental Material for details of this calculation). Although this type of analysis has a limited ability to precisely identify specific target genes, it can give us a general sense of whether PIWI/piRNA have an altered capacity to target and regulate gene expression in *piwi-MatKD* F1 embryos.

According to this analysis, approximately 50% of piRNA reads in all our libraries had the capacity to target at least one mRNA (Fig 5E). 63 genes had decreased piRNA targetability in *piwi-MatKD* compared to *GFP-MatKD* (Fig 5F, Table S5); that is, piRNAs antisense to these genes were abundant in *GFP-MatKD* control embryos, but were depleted in *piwi-MatKD* embryos. A few of these genes with changed piRNA targetability also had changed mRNA levels according to our RNA-seq. For example, the Juvenile Hormone Epoxide Hydrolase genes *Jheh2* and *Jheh3* had decreased piRNA targetability and increased mRNA expression (Fig 5G), indicating that their expression is normally repressed by Piwi and piRNAs. Juvenile Hormone impacts a myriad of processes in *Drosophila*, including longevity, stress response, and, notably, oogenesis (Santos et al. 2019). The *Jheh* gene family has not been studied extensively, but *Jheh2* is known to be expressed in male, but not female, embryonic gonads (Casper and Van Doren 2009). The observation that maternal Piwi and piRNAs can repress a gene that is specifically expressed in male PGCs led us to hypothesize that maternal Piwi could regulate other sex-biased germline genes, a mechanism that could explain the subfertility of *piwi-MatKD* F1 females.

### The *piwi-MatKD* F1 female germline is partially masculinized

Male-specific and female-specific characteristics of XY and XX PGCs, respectively, become evident around gonad coalescence (Casper and Van Doren 2009; Poirie et al. 1995; Staab et al. 1996; Wawersik et al. 2005), precisely the stage where we saw maternal Piwi protein was still maintained (Fig 1). In light of this and the fertility defects observed in *piwi-MatKD* F1 females, we asked whether female PGCs in *piwi-MatKD* F1 embryos were masculinized.

One key difference between male and female PGCs is the timing of their proliferation. PGCs form before cellularization of the embryo (Poirie et al. 1995). Normal numbers of PGCs were specified in *piwi-MatKD* F1 as compared to *GFP-MatKD* F1 embryos (Fig S9). After this initial phase, PGCs do not proliferate during migration to the embryonic gonad (Su et al. 1998), but male PGCs recommence proliferation at gonad coalescence, while female PGCs remain quiescent (Poirie et al. 1995; Wawersik et al. 2005). We saw this pattern reflected in *GFP-MatKD* F1 embryos: there were similar numbers of PGCs in male and female embryos at stage 15, just after gonad coalescence, but at stages 16 and 17, male PGCs increased in number while female PGCs did not (Fig 6A). In contrast, in female *piwi-MatKD* F1 embryos, PGC numbers increased at stages 16 and 17, resembling their sibling male embryos (Fig 6B-C). This change indicates that PGCs in female *piwi-MatKD* F1 embryos acquired a male PGC proliferation pattern.

**Figure 6.**
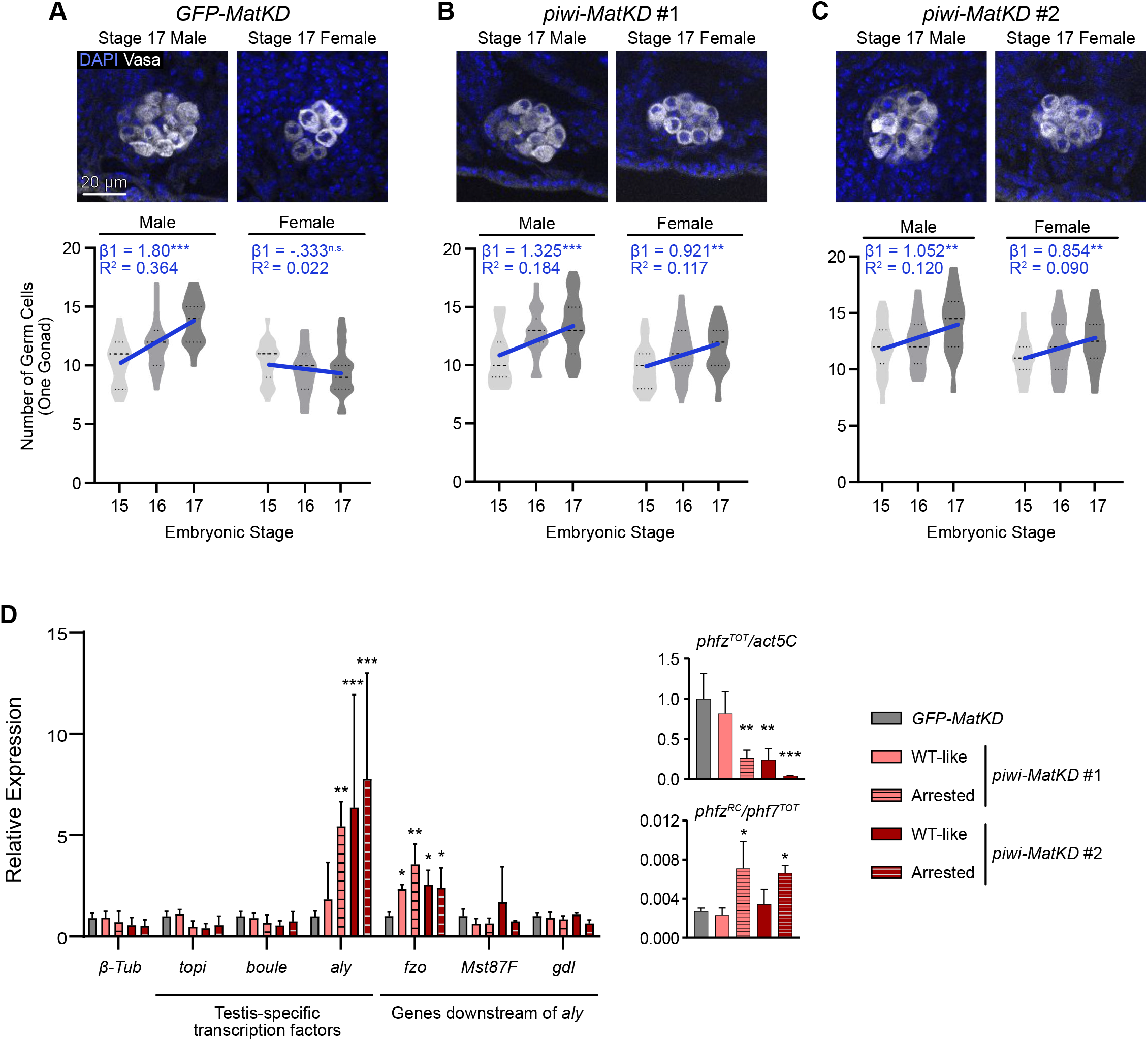
The *piwi-MatKD* F1 female germline is partially masculinized. (A - C) Germ cell numbers in male and female embryonic *MatKD* F1 gonads (stages 15-17) were visualized (upper panel) and counted (lower panels). Embryos were sexed by paternally inherited *Dfd-lacZ* on the X chromosome and embryos were staged based on gut morphology. Violin plots indicate germ cell count at each stage and condition; thick dashed line indicates median and thin dotted line indicates 1^st^ and 3^rd^ quartiles. Line indicates regression line of germ cell numbers of each sex within each genotype, β1 indicates the coefficient calculated from Poisson linear regression, and R^2^ indicates fit of the data to regression line. Asterisk indicates significance of the coefficient (** p < 0.01, *** p < 0.001) compared to the null hypothesis (β1 = 0). n = 66 - 81 per sex of each genotype, with 20-27 per stage of each sex of each genotype. (A) *GFP-MatKD* F1 embryos. (B) *piwi-MatKD* #1 F1 embryos. (C) *piwi-MatKD* #2 F1 embryos. (D) RT-qPCR for key testis-specific mRNAs in *MatKD* F1 ovaries, normalized to *actin5C* RNA levels (left panel). Upper right panel indicates expression level of total *phf7 (phf7^TOT^)* and lower right panel indicates *phf7^RC^* relative to *phf7^TOT^,* after each was normalized to *actin5C*. Two-way ANOVA and Dunnett’s Multiple Comparisons Test, * p<0.05, ** p<0.01, *** p<0.001.

To explore whether this apparent masculinization of the *piwi-MatKD* F1 female germline continued in the adult ovary, we used RT-qPCR to examine the expression of adult ovarian mRNAs. We did not see any consistent change in the expression of canonical ovary-specific genes in *piwi-MatKD* F1 ovaries (Fig S10). However, some canonical testis-specific genes were present at higher levels in *piwi-MatKD* F1 ovaries than in controls (Fig 6D). These genes included *always early* (*aly*), a core factor in spermatid differentiation (White-Cooper et al. 2000), and *fuzzy onion* (*fzo*), which is activated by *aly* during spermatogenesis (Chen et al. 2011) (Fig 6D, left panel). We also measured the expression of *phf7*, a histone reader important for spermatogenesis, and *phf7^RC^*, the male-specific isoform of *phf7* which causes oogenic arrest when overexpressed (Smolko et al. 2020). We observed that although the expression of *phf7* is overall decreased in *piwi-MatKD* F1 ovaries, the relative proportion of *phf7^RC^* was increased in *piwi-MatKD* F1 arrested ovaries as compared to *GFP-MatKD* F1 ovaries (Fig 6D, right panels).

The fact that only a subset of testis-specific genes we tested was derepressed in *piwi-MatKD* F1 ovaries is consistent with the fact that these ovaries do not have the classic tumor-like morphology of a dramatically masculinized female germline (Chau et al. 2009; Nagoshi et al. 1995; Steinmann-Zwicky et al. 1989) (Fig 3). Instead, we believe this represents a partial masculinization of the *piwi-MatKD* F1 ovary. We did not see defects in *piwi-MatKD* F1 somatic sex determination (Fig S11), so the partial masculinization in *piwi-MatKD* progeny appears to be germline-specific.

## DISCUSSION

In this paper, we have reported that maternal Piwi has long-ranging functions on the development of progeny. Maternal Piwi can be detected in primordial germ cells (PGCs) throughout embryogenesis, and in at least some third instar larval ovaries. Furthermore, our analyses revealed a novel function of maternal Piwi in ensuring the germline development of female progeny. Our transcriptome analysis, piRNA analysis, and examination of embryonic germ cell number indicate a partial masculinization of the *piwi-MatKD* F1 female germline that is known to cause oogenic defects and sterility (Chau et al. 2009; Nagoshi et al. 1995; Schupbach 1985; Steinmann-Zwicky et al. 1989; Yang et al. 2012). These findings reveal a novel sex-specific function of a maternal-effect gene.

### A novel female-specific role for a maternal factor

There have been decades of extensive study into maternal mutations that affect germline development in *Drosophila*, but most of these have focused on defects that manifest during early embryogenesis, such as the specification of PGCs (Schupbach and Wieschaus 1986; Seydoux and Braun 2006). Other studies have identified maternal-effect phenotypes that manifest in later PGC processes, such as migration through the primordial midgut (Asaoka-Taguchi et al. 1999; DeGennaro et al. 2011; Kunwar et al. 2003) and survival during migration (Kugler et al. 2013; Slaidina and Lehmann 2017). In the case of maternal Piwi, we observed its persistence in PGCs through gonad coalescence (Fig 1), implying that it can impact even late embryonic stages of germline development.

Indeed, we saw that maternal *piwi* knockdown caused defects in the embryonic coalesced gonads (Fig 6) as well as adult gonads (Fig 2, 3) of progeny. Strikingly, the defects in fertility and gonadal morphology in *piwi-MatKD* F1s were specific to females. This could reflect that the female germline has a more stringent requirement for Piwi levels than the male germline, which has previously been suggested for zygotically-expressed Piwi (Gonzalez et al. 2015), or that maternal Piwi is specifically required for the oogenesis of female progeny. Nevertheless, to our knowledge, no maternal factor has previously been shown to specifically affect the development of one sex but not the other.

Germline development of males and females is largely similar during embryogenesis until gonad coalescence, when sex-specific germline gene expression begins (Casper and Van Doren 2009) and male germ cells begin to proliferate while female germ cells remain quiescent (Poirie et al. 1995; Wawersik et al. 2005). Few studies, if any, have examined maternal-effect phenotypes at these late stages of embryogenesis, let alone in larval or adult stages. In light of our findings about maternal Piwi, it is likely that examining other maternal proteins for their longevity and functions beyond embryogenesis will reveal new developmental roles long after the maternal-to-zygotic transition.

### The embryonic PIWI/piRNA pathway and transposon control

Since early observations that maternal PIWI proteins and piRNAs accumulate in the germ plasm (Brennecke et al. 2007; Harris and Macdonald 2001; Megosh et al. 2006), it has been accepted that the maternal deposition of piRNAs shapes the piRNA pool going forward in development. So far, examination of this hypothesis has been focused on the molecular mechanisms of piRNA biogenesis and piRNA involvement in epigenetic regulation and transposon suppression (Akkouche et al. 2013; Brennecke et al. 2008; de Vanssay et al. 2012; Gu and Elgin 2013; Le Thomas et al. 2014a; Le Thomas et al. 2014b). In addition to directly binding and regulating zygotic genes, maternally deposited piRNAs can contribute to the generation of zygotic piRNAs via the secondary piRNA biogenesis pathway once zygotic mRNA substrates begin to be produced (Barckmann et al. 2018; Wang et al. 2015). For example, in *Bombyx mori*, maternally deposited piRNAs are stable for at least 12 hours during embryogenesis and trigger the generation of new piRNAs when they target zygotic mRNAs (Kawaoka et al. 2011). This means that loss of maternal piRNAs can affect gene expression beyond the maternal phase of embryogenesis. The best-characterized examples of this to date are when piRNAs targeting paternally-inherited transposons are absent in the maternal pool, resulting in transposon derepression and atrophic or arrested ovaries in adults (Akkouche et al. 2013; Brennecke et al. 2008). Our *piwi-MatKD* strategy resulted in a reduced pool of Piwi-bound piRNAs in the early embryo (Fig 5C), which could reflect either a disruption of piRNA biogenesis during oogenesis or instability of piRNAs unbound by their partner PIWI protein, or both. Regardless, this loss of maternal piRNAs almost certainly would change PIWI/piRNA target RNAs later in embryogenesis.

Despite the reduced pool of maternally deposited piRNAs, *piwi-MatKD* F1 individuals did not experience the dramatic derepression of transposons that is emblematic of disruptions to the PIWI/piRNA pathway (Fig 4). Although several transposons were consistently derepressed in *piwi-MatKD* F1 early embryos (Fig 4A), most of these still remained at fairly low expression levels (Fig S8). The exception was a group of telomere-associated transposons: *Het-A*, *TART,* and *TAHRE.* Derepression of these transposons during oogenesis can cause early embryonic chromosome segregation defects (Durdevic et al. 2018; Morgunova et al. 2015) that resemble defects of embryos strongly depleted of maternal Piwi (Mani et al. 2014), so the up-regulation of these transposons in *piwi-MatKD* #2 F1 embryos may partially explain their frequent embryonic arrest (Fig S3). However, it is unclear how the observed mild derepression of *Het-A, TART,* and *TAHRE* in the early embryo might relate to the female-specific *piwi-MatKD* F1 subfertility which manifested in adulthood, especially because these transposons were repressed in *piwi-MatKD* F1 adult ovaries (Fig 4D). If *Het-A*, *TAHRE*, and *TART* contributed to the *piwi-MatKD* F1 female subfertility, it must have been indirectly through transient activity.

Previous studies have shown that the PIWI/piRNA pathway is actively involved in transposon repression in gonads throughout development (Marie et al. 2017), and the transient knockdown of zygotic Piwi during mid- to late- embryogenesis (3-16 hours after egg laying) results in massive transposon derepression and subsequent sterility in adult ovaries (Akkouche et al. 2017). While the knockdown strategy used by Akkouche *et al.* reduced *piwi* mRNA levels during embryogenesis, it did not affect levels of maternal Piwi protein, which we have shown to remain present at high levels throughout embryogenesis (Fig 1). In contrast, because we saw that zygotic Piwi was barely detectable at mid-embryogenesis (Fig S2), any effect on zygotic *piwi* expression was likely negligible compared to the effect on maternal *piwi* expression in our knockdown strategy. Furthermore, our knockdown strategy reduced maternal Piwi protein levels starting from mid-oogenesis in F0 females (Fig 2A-B). As such, *piwi-MatKD* F1 embryos not only lacked the normal function of maternal Piwi during embryogenesis, but also inherited any dysregulation of gene expression (Fig 5A), piRNA biogenesis (Fig 5C-F), and possibly epigenetic state that occurred during oogenesis. Both of these embryonic Piwi knockdown strategies ultimately resulted in subfertile adult females, but the different effects on transposon repression in these two systems suggest that this subfertility is produced by different mechanisms. Akkouche *et al.* did not discuss the fertility of males depleted of zygotic Piwi. Given our report of female-specific germline defects upon maternal *piwi* knockdown, examining the males depleted of zygotic Piwi during embryogenesis will aid in the further disentanglement of maternal and zygotic Piwi functions during embryogenesis.

### The PIWI/piRNA pathway and sex determination

There is a growing understanding that the regulatory potential of the PIWI/piRNA pathway extends far beyond transposon suppression (Rojas-Rios et al. 2017; Wang and Lin 2021). This is especially well-understood in *C. elegans* and mice and is becoming increasingly apparent in *Drosophila* in the case of Aubergine (Barckmann et al. 2015; Ramat et al. 2020; Rouget et al. 2010; Vourekas et al. 2016). Given the extreme heterogeneity and low complementarity requirements of piRNAs (Barckmann et al. 2015; Gou et al. 2014; Halbach et al. 2020; Shen et al. 2018; Zhang et al. 2018), it is not surprising that they can target and regulate a wide variety of RNAs.

In the context of this study, perhaps the most relevant examples of PIWI proteins regulating non-transposon mRNAs are in the cases of sex determination in *B. mori* and *C. elegans.* Despite otherwise very different modes of sex determination, individual piRNAs have been implicated in this process for both organisms. In *B. mori,* the *Fem* piRNA targets the mRNA of male sex determination gene *Masc* for degradation in females (Kiuchi et al. 2014), while in *C. elegans*, the *21ux-1* piRNA targets the mRNA of male sex determination gene *xol-1* for degradation in hermaphrodies (Tang et al. 2018). When these piRNAs were deleted, the organisms became partially masculinized. Although we did not identify specific piRNAs involved in sex determination in *Drosophila* in this study, the partial masculinization of *piwi-MatKD* F1 female PGCs and ovaries (Fig 6) suggests a role for the *Drosophila* PIWI/piRNA pathway in repressing male gene expression programs in the female germline.

Both in *C. elegans* and in our study, modulation of PIWI proteins and/or piRNAs did not fully transform XX individuals into males, but rather moderately masculinized them (Tang et al. 2018) (Fig 6), implying a partial shift from female gene expression patterns to male gene expression patterns. This emerging link between the PIWI/piRNA pathway and sex determination may be yet another illustration of how piRNAs can be cellular tools to “tune,” rather than completely switch, gene expression programs.

## Supporting information

Supplemental Figures and Scripts

Reagents Table

## ACKNOWLEDGEMENTS

We thank the Developmental Studies Hybridoma Bank for providing antibodies, and the Bloomington Drosophila Stock Center, Drs. Ting Xie, Trudi Schupbach, and Mark van Doren for providing fly stocks. We thank Drs. Mark van Doren, Nils Neuenkirchen, Hongying Qi, Valerie Reinke, Lynn Cooley, and Karla Neugebauer for discussion and technical support, and Drs. Celina Juliano, Nils Neuenkirchen, Hongying Qi, and Valerie Reinke for valuable comments on the manuscript. Dr. Nils Neuenkirchen also wrote our custom script (available in Supplemental Materials) for adding an NH tag to Bowtie output files in the piRNA sequencing analysis. Sequencing service was conducted at the Yale Stem Cell Center Genomics Core facility, which was supported by the Connecticut Regenerative Medicine Research Fund and the Li Ka Shing Foundation.

## Author Contributions

LG and HL conceived and designed the project. LG conducted all experiments and analysis, except XT conducted and analyzed larval gonad immunofluorescence experiments in Fig S1. LG prepared the figures and the initial draft. LG, HL, and XT revised the paper. HL supervised the study.

