## Supplemental Figures and Scripts for "Maternal Piwi Regulates Primordial Germ Cell Development to Ensure the Fertility of Female Progeny in *Drosophila*"

### Supplemental Materials

TABLE OF CONTENTS


| | |
|---|---|
| Supplemental Figures 1-11 | pp 2-11 |
| Supplemental Methods (Scripts 1-3) | pp 12-15 |

### SUPPLEMENTAL FIGURES

#### Supplemental Figure 1

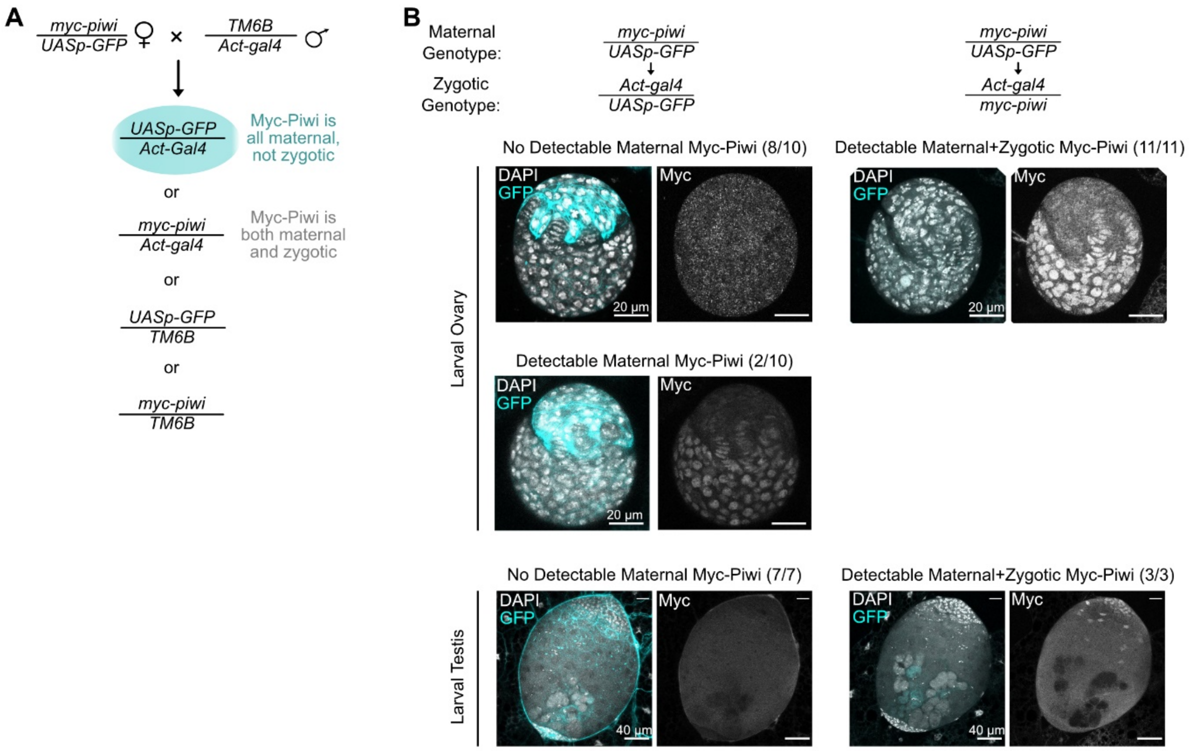


**Figure S1. Maternal Piwi protein is detectable in a minority of L3 ovaries.** (A) Schematic of crossing strategy to visualize maternal Piwi. (B) Representative images of immunostaining for maternal Myc-Piwi in larval ovaries and larval testes from the cross depicted in Fig. S1A. Numbers in parentheses are (# individuals with indicated Myc-Piwi expression pattern / total # individuals of indicated genotype).

**Supplemental Figure 2**

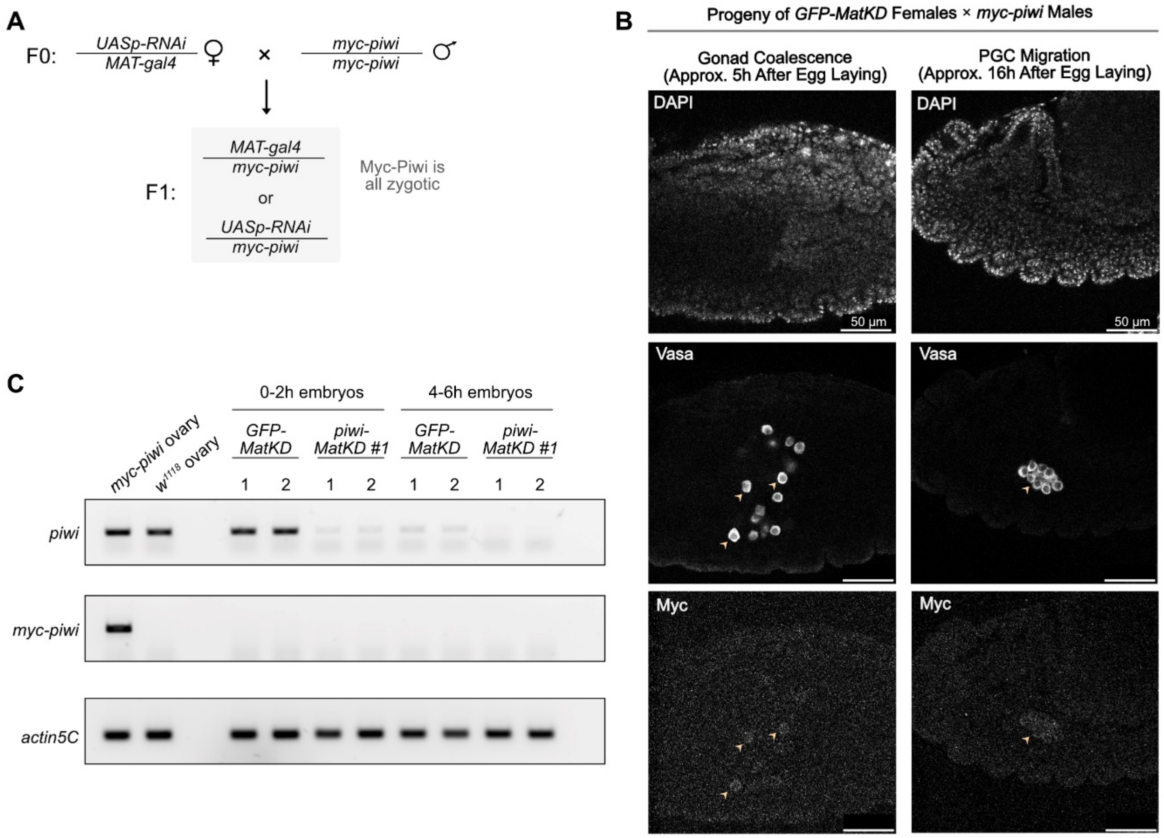


**Figure S2. Zygotic Piwi is expressed at nearly undetectable levels in mid-embryogenesis.** (A) Schematic of crossing strategy to visualize zygotic Piwi (from paternally-inherited *myc-piwi*). (B) Representative images of immunostaining for zygotic Myc-Piwi from control (*GFP*-MatKD) mothers. Zygotic Myc-Piwi is not quite detectable. (C) RT-PCR for total *piwi*, *myc-piwi*, and *actin5C*. Left two lanes are positive and negative controls, respectively, for *myc-piwi* expression, and remaining lanes are from embryos collected from the cross depicted in Fig S2. Only embryos laid by *piwi-MatKD* #1 mothers were used because embryos laid by *piwi-MatKD* #2 mothers are mostly arrested in early embryogenesis (see Fig S3), precluding analysis of RNA levels in bulk embryos beyond the 0-2h stage.

**Supplemental Figure 3**

A

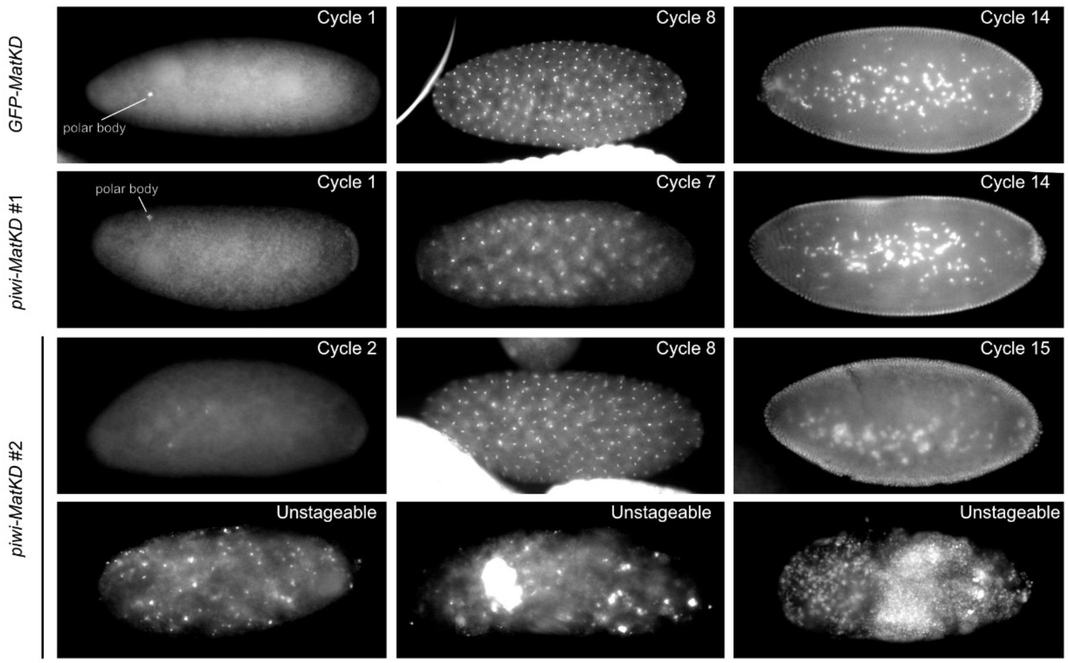


B

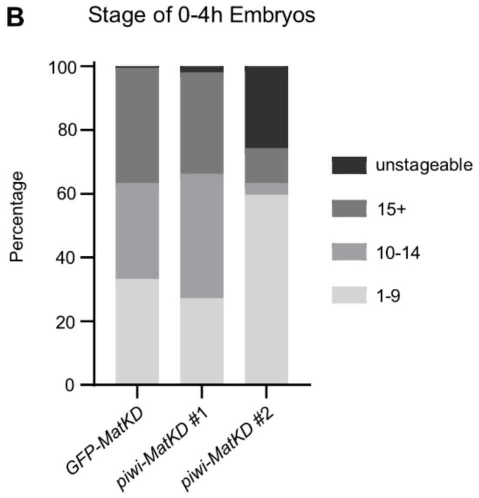


**Figure S3. *piwi-MatKD* #2 F1 embryos suffer a high rate of embryonic arrest.** (A) Representative images of 0-4h F1 embryos laid by *GFP-MatKD* and *piwi-MatKD* females. Images are at the same scale. (B) Relative frequency of F1 embryos at these stages. n=109-199.

**Supplemental Figure 4**

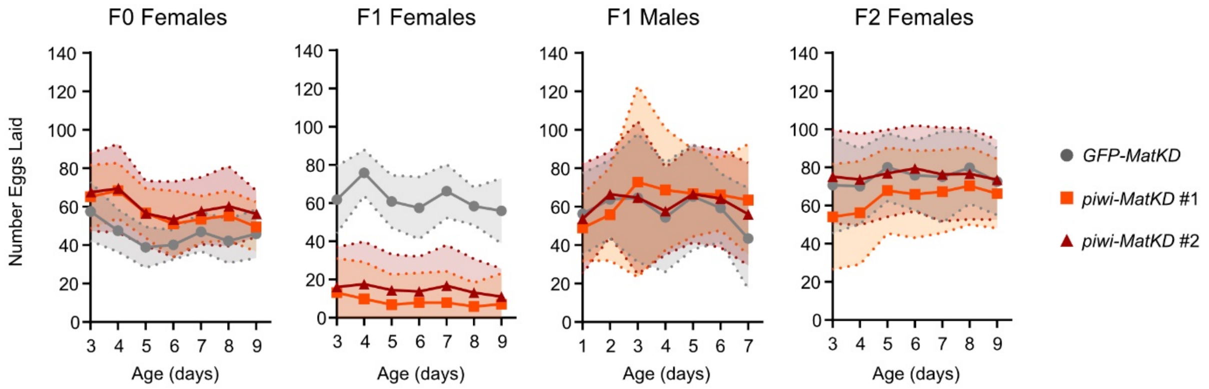


**Figure S4. Number of eggs laid daily by *GFP-MatKD* or *piwi-MatKD* F0, F1, and F2 individuals.** 7-day fertility tests of individual females mated to two $w^{1118}$ males or individual males mated to three $w^{1118}$ females (see cross in Fig. 2A). These data are also represented in total embryo counts in Fig 2D-G, upper panels. Mean ± SD, n=16-24.

**Supplemental Figure 5**

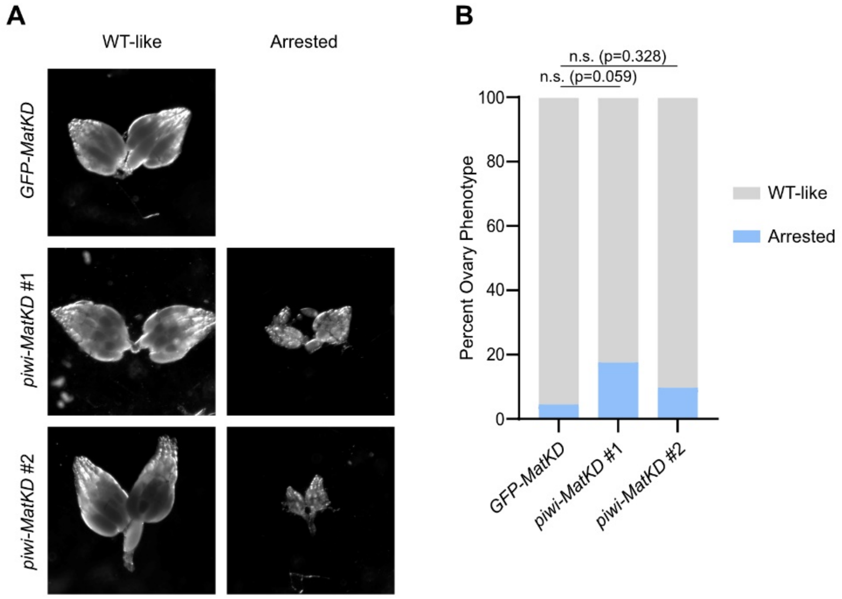


**Figure S5*. piwi-MatKD* F2 ovaries have largely normal morphology**. (A) Representative images of ovaries from *MatKD* F2 females (see cross in Fig 1A). Ovary images are at the same scale. (B) Quantification of the relative frequency of WT-like and arrested ovaries from *MatKD* F2 females. No Agametic ovaries were observed. n=34-51 per genotype, Chi-square Test.

**Supplemental Figure 6**

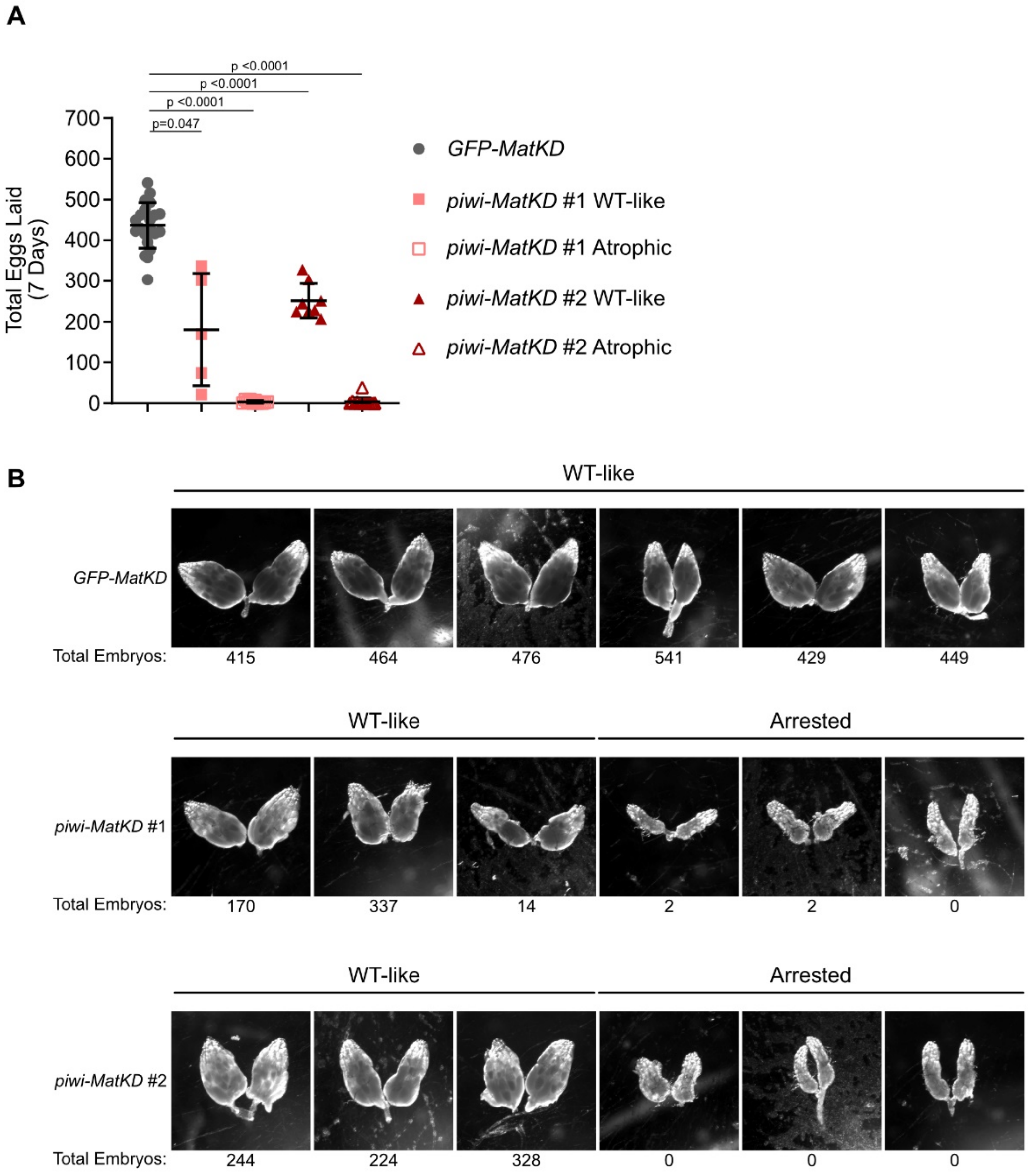


**Figure S6. *piwi-MatKD* F1 females with both WT-like and arrested ovaries are subfertile compared to *GFP-MatKD* F1.** (A) Total eggs laid over seven-day fertility test by individual females of the indicated genotype and ovary morphology. One-Way ANOVA and Dunnett's Multiple Comparisons Test. n=5-23. (B) Representative images of *MatKD* F1 ovaries, all taken at the same magnification. The number below each image is the total number of eggs laid by that individual during the seven-day fertility test. Ovary images are at the same scale.

**Supplemental Figure 7**

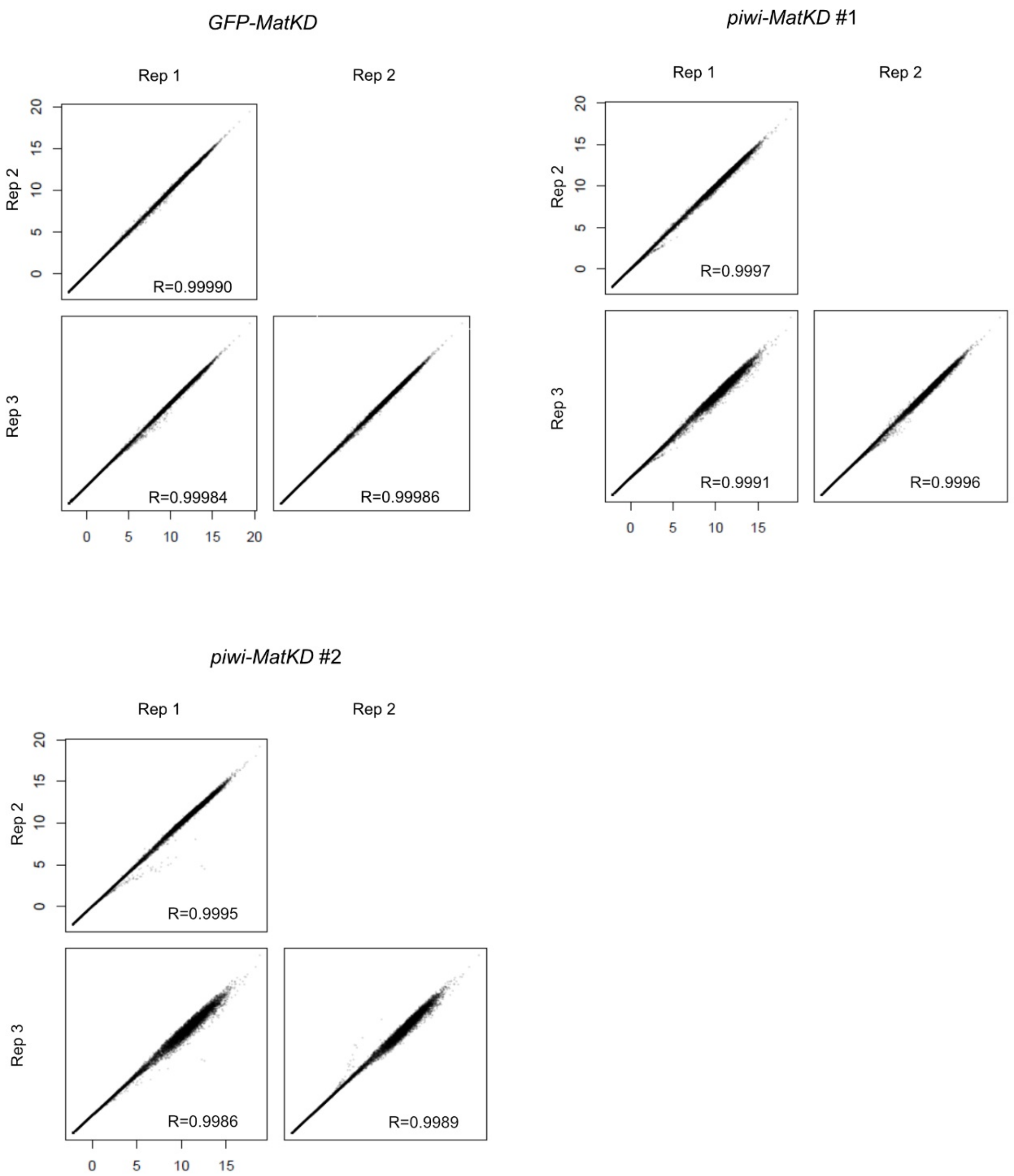


**Figure S7. RNA-seq data from *MatKD* F1 embryos within each genotype are highly reproducible.** After gene expression levels were determined using TETools and DESeq2, pairwise comparisons were made among the regularized log transformed read counts from three replicates of each genotype, and are presented as scatterplots. R indicates correlation coefficient.

**Supplemental Figure 8**

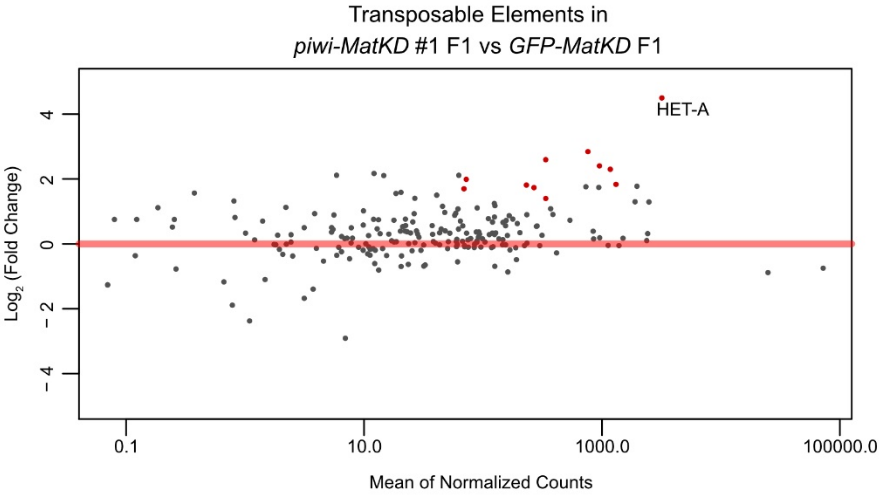


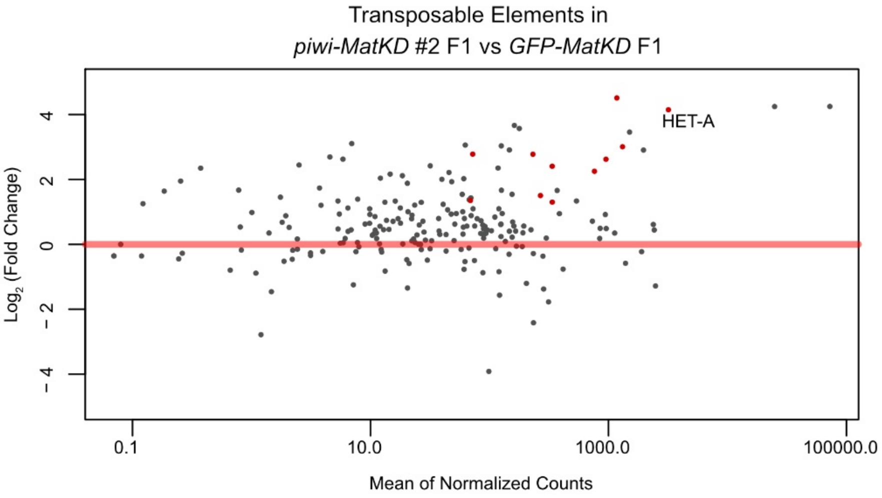


**Figure S8. Most transposons that were consistently derepressed in the *piwi-MatKD* F1 early embryo were present at low levels.** MA plots of mean normalized read counts for transposons in *piwi-MatKD* vs *GFP-MatKD* early embryos. Red points indicate transposons which were consistently derepressed (see Fig. 4).

**Supplemental Figure 9**

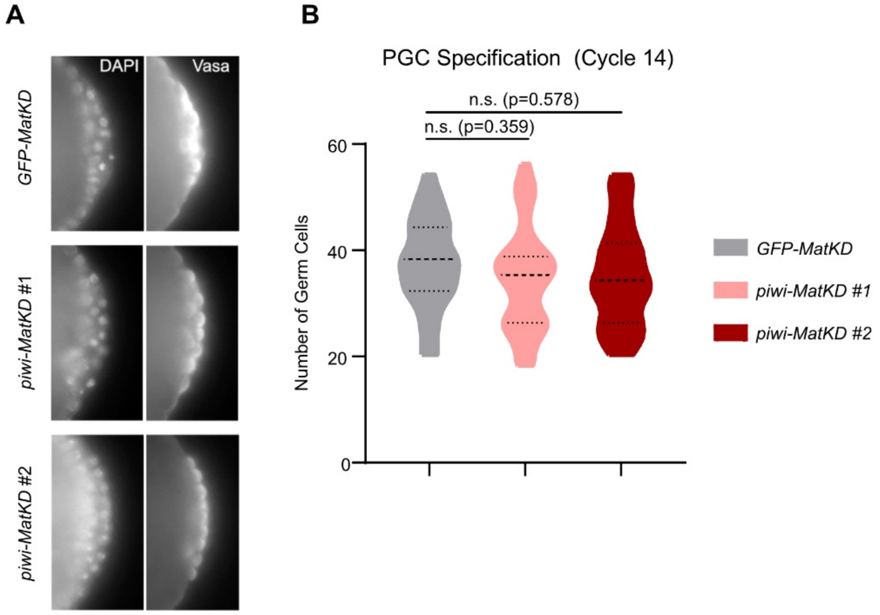


**Figure S9. Number of PGCs specified in *piwi-MatKD* F1 embryos does not differ from that in *GFP-MatKD* F1 embryos.** (A) Representative images of Cycle 14 embryos. Images are at the same scale. (B) Quantification of number of Vasa-positive cells at the posterior pole of Cycle 14 embryos. One-Way ANOVA and Dunnett's Multiple Comparison's Test. n=29-38.

**Supplemental Figure 10**

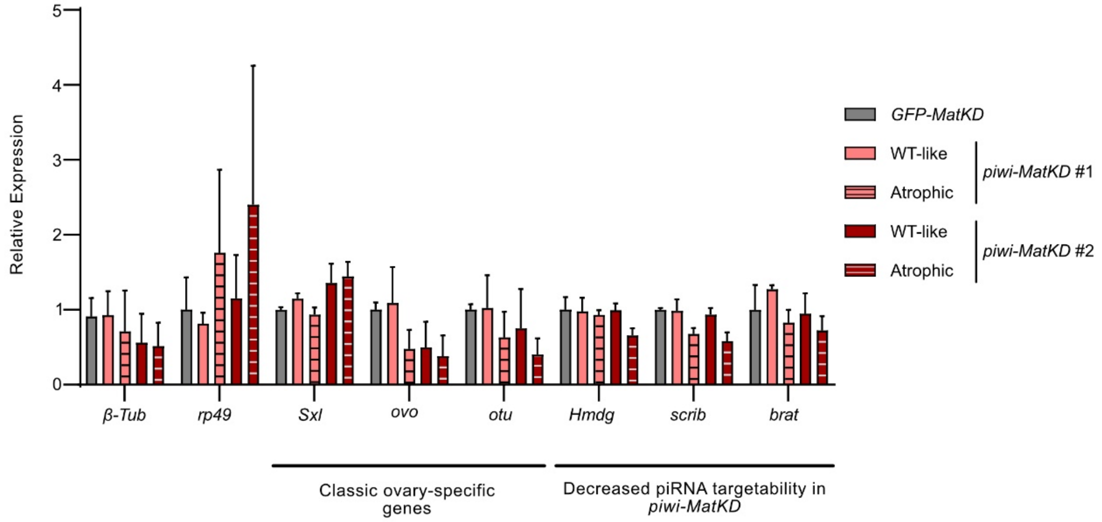


**Figure S10. The expression of ovary-specific genes in *piwi-MatKD* F1 ovaries is unchanged.** RT-qPCR for ovary-specific mRNAs in *MatKD* F1 ovaries. Expression was normalized to *actin5C* RNA levels. All differences in gene expression were nonsignificant.

**Supplemental Figure 11**

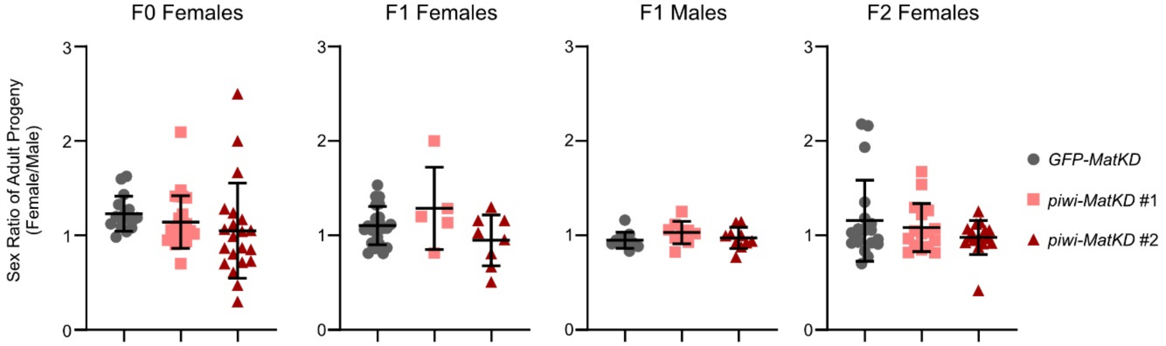


**Figure S11. Sex ratios of adult progeny from *MatKD* F0, F1, and F2 parents.** Female-to-male ratios of adult progeny from the indicated parents mated to 2-3 $w^{1118}$ flies of the reciprocal sex (see cross in Fig 2A. These adults are also quantified without distinguishing between males and females in Fig 2D-G, lower panels). Sex ratios were only calculated for crosses with ≥10 adult progeny. All differences in sex ratios were nonsignificant.

**SUPPLEMENTAL METHODS**

**Script 1: Create list of putative piRNA sequences from total Small RNA-seq**

```
module load Bowtie/1.2.2

#map to rRNA, tRNA, siRNA
#remaining sequences are miRNAs and piRNAs

bowtie …/genomes/dm6/ncrna -v 1 -a 41550_rep1_R1.trimfilt.fastq.gz
41550_rep1_R1_ncrna.map --un 41550_rep1_R1_mipirna.map

#map to hairpin (miRNAs)
#remaining sequences are putative piRNAs

bowtie …/genomes/dm6/hairpin -v 1 -a 41550_rep1_R1_mipirna.map
41550_rep1_R1_hairpin.map --un 41550_rep1_R1_nomirna.fa

#map to genome
#only require one mapping site, can be multimapping

bowtie …/genomes/dm6/genome -S -n 1 -k 1
41550_rep1_R1_nomirna.fa>41550_rep1.pirna.sam
```

**Script 2: piRNA target analysis**

```
module load Bowtie/1.2.2
module load GitPython/2.1.11-foss-2018b-Python-3.7.0
module load Subread/2.0.0-GCC-7.3.0-2.30

#aligns piRNAs to the transcriptome: antisense, allowing 3 mismatches in first 24 nucleotides,
report all targets
#"41550_rep1_pirna.fq" is the list of piRNAs that align anywhere in the genome for that replicate
#this will find all targets (-a) that are reverse-complementary to genes (--nofw)
#with up to 3 mismatches (-v 2) in the first 24 nucleotides (-l 24)

bowtie …/dm6_genes -v 2 -a --nofw -l 24 -S …/41550_rep1_pirna.fq.gz
>41550_rep1_pirna_targets.sam

#adding "NH" tag to all alignment records in the SAM file
#so featureCounts can assign weighted counts based on the number of times a particular
piRNA read aligned within the transcriptome

python3 …/NH_insert_v2.py 41550_rep1_pirna_targets.sam

#counting reads at each feature
#-M says count multi-mapping reads
```

#--fraction says weight those reads based on how many sites they map to (so weight as 1/x, where x is the number of mapping locations)
#-F GTF specifies that it'll take an GTF file
#-a specifies the annotation file (GTF)
#-o specifies the output file

featureCounts -M --fraction -F GTF -a dmel.GTF.txt -o 41550_rep1.wtargetsfrac.counts 41550_rep1_pirna_targets_NH_v2.sam

**Script 3: Python script ("NH_insert_v2.py" above) for calculating and adding NH flag, to indicate the number of alignments for each piRNA read.**

```
#!/usr/bin/env python3

import time
t0 = time.perf_counter()

import sys
input = sys.argv
input_len = len(input)
input_filename = ''
output_filename = ''

print('''
----------------------------------------------------------------------
XM:i:(n) to NH:i:(n-1) conversion,                          ver1.2

written by Nils Neuenkirchen, PhD                          04/02/2020
----------------------------------------------------------------------
Command line entry:
python NH_insert.py FILENAME
----------------------------------------------------------------------
Input example:
K00175:212:HC2KCBBXY:2:2228:29376:49230 16      FBgn0250816     57306   255     25M     *
0       0       ACTTCAGCACTCATCTCATTAATAA       JJJJJJJJJJJJJJJJJJJJFFFAA       XA:i:1
MD:Z:18G6       NM:i:1  XM:i:7

Output example:
K00175:212:HC2KCBBXY:2:2228:29376:49230 16      FBgn0250816     57306   255     25M     *
0       0       ACTTCAGCACTCATCTCATTAATAA       JJJJJJJJJJJJJJJJJJJJFFFAA       XA:i:1
MD:Z:18G6       NM:i:1  XM:i:7  NH:i:6
----------------------------------------------------------------------''')

# Checking input file in command line
if input_len == 2:
   input_filename = str(input[1])
   print('Loading file: ' + input_filename)
   output_filename = input_filename[:-4] + '_NH_v2' + input_filename[-4:]
   print('Output file:  ' + output_filename)
```

```
    print('-----------------------------------------------------------------------')
else:
    print("Please enter file name\npython NH_insert.py FILENAME")
    print('-----------------------------------------------------------------------')
    sys.exit() # exits script immediately if too few or to many arguments are given


# Fast approach
# Reads file, processes each line, and immediately writes it into new file
# 1) reads every line
# 2) only if the line contains XM:i:, and additional value of NH:i:(n-1) is added
# 3) any other lines are just returns are is

output = open(output_filename, 'w')

with open(input[1]) as my_file:
    for line in my_file:
        if line.count('XM:i:') == 1:
            output.write(line[:-1] + '\tNH:i:' + str(int(line[line.find('XM:i:')+5:-1])-1) + '\n')
        else:
            output.write(line)
output.close()

print('\nDone...')

t1 = time.perf_counter()
print(f'Running time: {t1-t0} seconds')
```
